# Revival of traditional agricultural systems – A multidisciplinary on-farm survey of maize-bean intercropping reveals unexpected competition effects on beans

**DOI:** 10.1101/2024.04.24.590929

**Authors:** Noa Vazeux-Blumental, Laura Mathieu, Théo Trabac, Carine Palaffre, Bernard Lagardère, Maryse Carraretto, Cyril Bauland, Martine Le Guilloux, Christine Paysant Le Roux, José Caïus, Anne Marmagne, Jérôme Enjalbert, Timothée Flutre, Edith Le Cadre, Virginie Parnaudeau, Daniel Muller, Yvan Moënne-Loccoz, Domenica Manicacci, Maud I. Tenaillon

## Abstract

Cereal-legume intercropping is a promising strategy for sustainable agroecosystems. The traditional intercropping of maize and bean is experiencing a revival in some modern agricultural settings, such as in southwestern France, where maize hybrids are intercropped with the commercialized Tarbais bean. We conducted on-farm surveys and a field assay to address the following questions: How does the cropping system impact yield, nutrient uptake, and rhizosphere bacterial assemblages? Do positive or negative interactions between maize and beans dominate in intercropping? What is the effect of intercropping on plant transcriptomics?
We recorded farming practices, conducted yield and nutrient measurements, and characterized soil bacterial assemblages to compare sole-cropped maize and beans with intercropped plants. A controlled field assay was also established to extend this comparison to plant gene expression differences.
Intercropping was associated with a trend towards increased bacterial diversity. The cropping system significantly influenced agronomic traits, with frequent farm-by-cropping system interactions underscoring the critical role of farming practices. Competition dominated maize-bean intercropping, with 34 negative correlations among the 47 significant ones between maize and bean traits. This competition affected yield and nutrition, but primarily impacted beans, which produced fewer but bigger/heavier seeds. Transcriptomic results concurred with these findings, revealing no differentially expressed genes in maize but 5,070 in beans under competition.
Overall, our findings suggest that beneficial interactions between the two crops are hindered under current field conditions, underscoring the importance of carefully considering partner varieties and farming practices to revive traditional agricultural systems.

**Societal Impact Statement:** Cereal-legume intercropping is a promising strategy for sustainable agroecosystems that leverages the biological complementarities between plant species, reducing the need for chemical inputs while enhancing field biodiversity. Here, we focused on maize-bean intercropping, which is experiencing a revival in modern conventional agricultural settings. We found a trend towards an increased soil bacterial diversity in intercropping. However, competition between the two crops dominated and primarily impacted yield and gene expression in beans. Despite lower yield, bean seeds exhibited greater size. Our study indicates that implementing variety selection of both species and adapting farming practices is essential to fully harness the potential of intercropping.

## 1. INTRODUCTION

Conventional agriculture mainly relies on simplified cropping systems, with a single crop species grown per field (sole-cropping) and environmental uniformization through chemical inputs to ensure high yields. Given the current environmental challenges, there is a growing concern for more sustainable agroecosystems based on ecological regulations (Beillouin 2021; Fréville et al. 2022; Gaba et al. 2015). In this context, mixed intercropping (hereafter termed intercropping), where several crops are grown simultaneously with no distinct row arrangements, relies on biological complementarities and beneficial species interactions that diminish input requirements (Bedoussac et al. 2015). This is the case for cereal-legume intercrops whose success depends on their complementarity, which results primarily from the capacity of legumes (*Fabaceae*) to assimilate nitrogen fixed by symbiotic rhizobia. By mixing cereals (*Poaceae*) with legumes, the need for nitrogen-based fertilizers is reduced when compared to sole-cropped cereals. Cereals promote symbiotic nitrogen fixation by depleting soil nitrogen accessible for legumes and also through their root exudation, stimulating nodulation and rhizobia symbiotic activity (Rodriguez 2020; Li et al. 2016a). Cereal-legume interactions can yield additional benefits. First, improved resource utilization may occur through the complementarity of their aerial and root architectures. This has been demonstrated in maize-bean-squash intercropping (Zhang et al. 2014), where root complementarity increases biomass production in N-deficient soils. Second, facilitative processes involving soil acidification by legume root exudates, which locally increases the availability of inorganic phosphorus in the soil, may benefit the cereal crop. This, for example, is exemplified in maize-faba bean (Li et al. 2007; Zhang et al. 2016), maize-chickpea (Li et al. 2004), wheat-faba bean (Li et al. 2016b), and wheat-common bean (Li et al. 2008) intercropping. Third, weed reduction can result from enhanced competitive abilities of the crop mixture, which decreases the need for herbicides (Poggio 2005). Fourth, the transfer of N and P may be accrued between plant species interconnected through arbuscular mycorrhizal fungi (Johansen and Jensen 1996; Zhang et al. 2019). Fifth, the recruitment of plant growth-promoting rhizobacteria, such as phloroglucinol-producing *Pseudomonas* on cereal roots, can promote plant growth and control over various root pathogens (Mazzola 2004).

Although beneficial interactions between cereals and legumes are well-documented from an agronomic perspective, the genetic and molecular determinants of their interactions and their impact on soil ecology remain poorly explored (Becker et al. 2023). This is particularly true for the intercropping of maize with climbing common bean, where the maize stalk supports the climbing bean, promoting its exposure to light. Yet this association is of significant interest for both economic and historical reasons. Economically, maize is the second most widely grown crop in the world, and the common bean is the most commonly used legume for direct human consumption. Historically, both crops were domesticated in Mexico (Bitocchi et al. 2013; Matsuoka et al. 2002), and their association within the same cropping system dates back to 7,000 to 4,400 years BP (Zizumbo-Villarreal and Colunga-GarcíaMarín 2010). This cropping system, central to the iconic “milpa”, featured maize, squash, and beans—collectively known as the “three sisters”—and was the predominant multi-cropping subsistence system in Mesoamerica. Notably, the practice of intercropping maize and beans has spread beyond its region of origin, such as on small farms in sub-Saharan Africa (Nassary et al. 2020) and in various traditional European farming systems (Vazeux-Blumental et al. 2024). Thus, the intercropping of maize and bean appears to have been introduced and maintained as these crops spread beyond their centers of domestication.

While being abandoned in favor of sole-cropping during the 20^th^ century, this practice has persisted on small-scale subsistence farms in many parts of the world and has recently experienced a revival in some modern agricultural settings. An example is the Tarbais bean in southwestern France, where maize and beans were introduced in the 18th century and were widely intercropped, becoming an essential part of the culinary heritage (Bonnain-Dulon and Brochot 2004). However, during the 1950s, the cultivation of beans considerably declined due to the intensification of agriculture and the introduction of maize hybrids i.e., maize varieties mainly used for animal feed and developed through crosses of inbred lines, significantly increasing yield and supporting industrial farming. As beans were unsuitable for mechanization, their cultivation around Tarbes (Hautes-Pyrénées area) decreased from approximately 12,000 hectares at the beginning of the 20th century to around 55 hectares by 1970 (Bonnain-Dulon and Brochot 2004). It was not until 1986 that the Tarbais bean was developed to diversify local agriculture and enhance farming profitability. This development promoted a local crop used in traditional dishes, such as the cassoulet stew. Tarbais bean obtained a quality label and a protected geographical indication in 2000, which makes it an emblematic crop in the region with high added value. Along with the increase in acreages of Tarbais beans, several farmers reintroduced intercropping with maize as an alternative to the use of plastic nets in sole-cropped beans (Figure 1).

**FIGURE 1.**
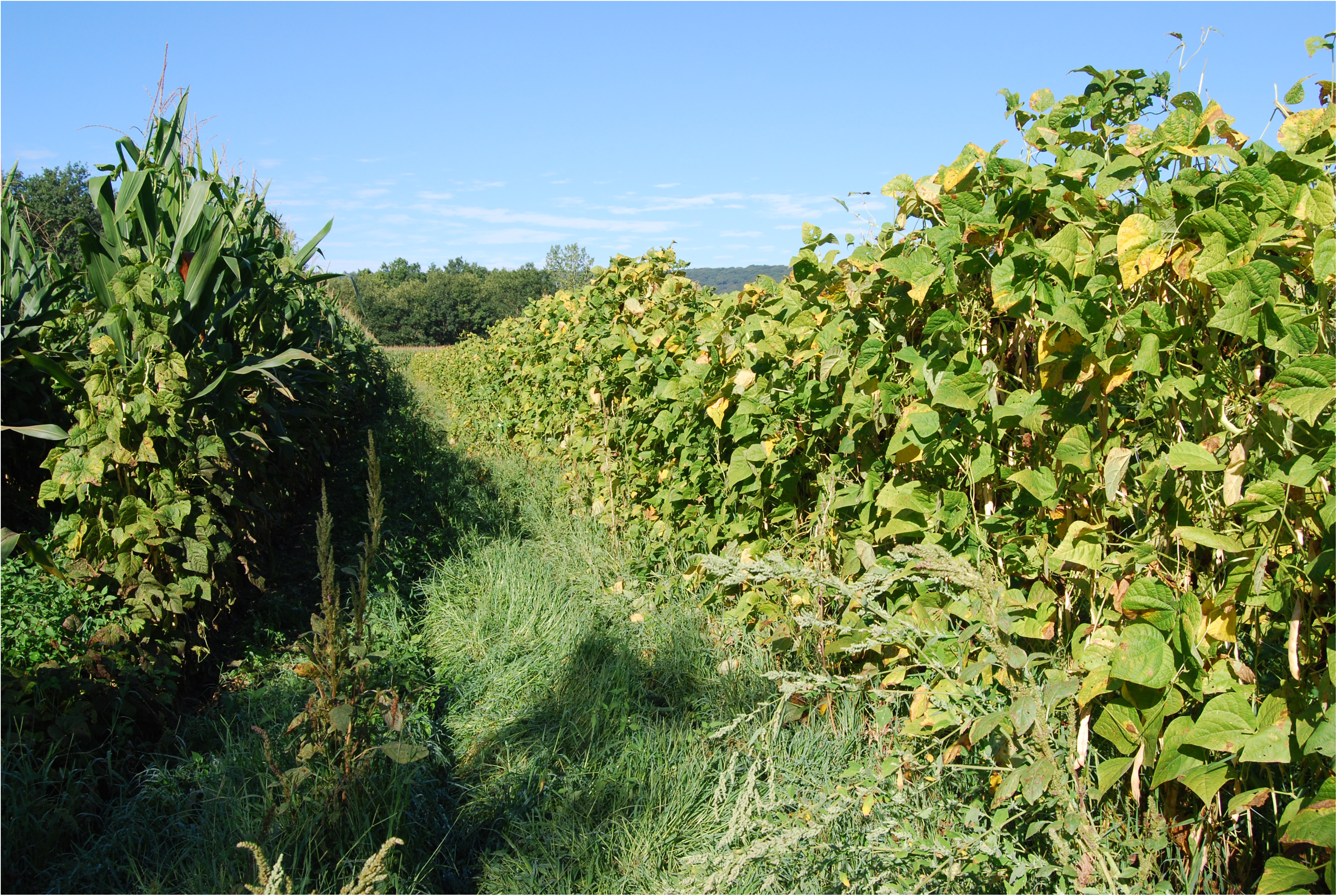
The local Tarbais bean intercropped with maize (left) or as a sole crop (right).

The objective of our study was to gain a deeper understanding of the functioning and potential benefits of intercropping maize and common beans. We focused on the Tarbais-maize system, which represents a revival of a traditional farming system using modern varieties and, most often, conventional farming practices (notably relatively high levels of nitrogen fertilization). While these practices may not be ideal for leveraging beneficial interactions between crops (Fréville et al. 2022), we hypothesized that maize-bean intercropping could still offer benefits. Due to the lack of research on this particular intercropping, a descriptive multidisciplinary approach combining agronomy, microbial ecology, and transcriptomics was adopted to answer three main questions: How does the cropping system impact yield, nutrient uptake and rhizosphere bacterial assemblages? Do positive or negative interactions between maize and bean dominate in intercropping at the phenotypic level? What is the effect of intercropping at the plant transcriptomic level? We recorded farming practices, performed agronomic and nutrient measurements on plants collected from intercropped and sole-cropped plots in farmers’ fields, and examined bacterial communities and corresponding soils in the rhizosphere. Because the detection of molecular signatures of intercropping requires a controlled environment with replicates, a we also established a controlled field assay in which both agronomic traits and gene expression were monitored to compare sole-cropping and intercropping of Tarbais bean with two maize hybrids.

## 2. MATERIALS AND METHODS

### 2.1 On-farm analyses in the Tarbes region

#### 2.1.1 Farms and farming practices

In the Tarbes region (Figure S1), located in southwestern France, farms that practiced maize and/or bean sole-cropping in addition to intercropping were identified, seven of which agreed to participate in this study. In this region, most farmers use the recent “Label Rouge” bean variety Alaric, an inbred line derived from crosses among local landraces registered in the European Community’s varietal catalog in 1999. The Lapujole variety, registered in 2014, was derived from Alaric by the introgression of an anthracnose resistance allele from the scarlet runner bean (*Phaseolus coccineus*). Therefore, both varieties are genetically similar (hereafter, designated Tarbais bean). Five of the seven farmers in our study used the Tarbais bean, and two used local bean landraces of their own. Most maize varieties were modern hybrids (Dataset S1). A single farmer (F4) used a local maize landrace. Only two farms (F6 and F7) practiced the three cropping systems (sole-cropped bean, sole-cropped maize and maize-bean intercropping). Farming practices in each farm were recorded (Dataset S1).

#### 2.1.2 Plant and soil analyses

All plant and soil samples were collected before harvest in October 2017, from 16 fields (1 to 3 fields per farm, Dataset S1). In each field, three randomly sampled plots (located > 5 m apart) served as replicates in the analyses. Plots comprised 4 to 5 (sole-cropping) or 7 to 10 (intercropping) plants each. Among the 16 fields, 7 (i.e., 21 plots) corresponded to intercropping, 5 (i.e., 15 plots) to maize sole-cropping, and 4 (i.e., 12 plots) to bean sole-cropping. Since maize and beans were collected separately in intercropping plots, these 48 plots gave 69 plant samples of 4-5 pooled plants each (except 1 plot with 3 plants). In addition, we retrieved about 500 g of soil from each plot (5-15 cm depth) to determine soil properties.

##### Soil analyses

Soil analyses were performed at the Laboratoire des sols d’Arras (INRAE), and 13 variables were determined, following standard procedures (see Dataset S2).

##### Plant phenotyping

All analyses were done on pooled plants from the same plot, targeting 22 maize traits and 24 bean traits (Dataset S4, Table 1) after separating vegetative (including maize cobs and bean pod shells) from reproductive (kernels for maize and seeds for beans, referred to as “propagules” in Table 1) parts. Propagules were counted per plant, but also per ear for maize and per pod for bean (referred to as “sets” in Table 1). Samples were finely ground, and subsamples (0.8–1.2 mg) were used for total N and C contents and ^15^N abundance, using the FLASH 2000 Organic Elemental Analyzer coupled to an isotope ratio mass spectrometer (ThermoFisher Scientific, Waltham, MA). The N and C concentrations (N% and C%) of vegetative parts (NV, CV) and propagules (NP, CP) were multiplied by dry weight per plant to compute N and C contents per plant for vegetative parts (NV_p, CV_p) and propagules (NP_p, CP_p).

**TABLE 1.** Description of phenotypes and nutrient variables measured on plants. ^a^: only measured for bean.

| Variable names | Variable description |
| --- | --- |
| Plants | Number of Plants per sample |
| Vw | Vegetative parts dry weight per plant (g) |
| Pw | Propagule dry weight per plant (g) |
| NbP | Number of Propagules produced per plant |
| Ple | Average Propagule length – measured on 50 propagules (mm) |
| Pwi | Average Propagule width – measured on 50 propagules (mm) |
| Psu | Average Propagule surface – measured on 50 propagules (mm<sup>2</sup>) |
| *d15NP* | Delta <sup>15</sup>N among total N of the Propagules |
| *d13CP* | Delta <sup>13</sup>C among total C of the Propagules |
| *d15NV* | Delta <sup>15</sup>N among total N of the Vegetative parts |
| *d13CV* | Delta <sup>13</sup>C among total C of the Vegetative parts |
| NP | Percent Nitrogen (measured in dry matter) in Propagules |
| CP | Percent Carbon (measured in dry matter) in Propagules |
| NV | Percent Nitrogen (measured in dry matter) in Vegetative parts |
| CV | Percent Carbon (measured in dry matter) in Vegetative parts |
| NP_p | Per plant Nitrogen content (g) in Propagules |
| CP_p | Per plant Carbon content (g) in Propagules |
| NV_p | Per plant Nitrogen content (g) in Vegetative parts |
| CV_p | Per plant Carbon content (g) in Vegetative parts |
| NbS | Number of Sets of propagules (ears or pods) per plant |
| PS (=NbP/NbS) | Average number of Propagules per Set – ears or pods |
| Sw (=Pw/NbS) | Average propagule weight per Set – ears or pods (g) |
| TPw (=1000×Pw/NbP) | Thousand Propagules dry weight (g) |
| *d15NVdfa*<sup>a</sup> | Nitrogen in the Vegetative parts derived from symbiotic fixation |
| *d15NPdfa*<sup>a</sup> | Nitrogen in the Propagules derived from symbiotic fixation |

The mineralization of soil nitrogen (kg·ha^-1^) was estimated in the plots using Nsim (Parnaudeau et al. 2012), based on soil characteristics (i.e., texture) (Clivot 2019), soil water content and daily temperatures recorded in the Tarbais region in 2017. Soil water content depends on soil characteristics, weather, and canopy conditions. Input parameters were calendar dates of sowing and harvest and the irrigation and nitrogen supply (Dataset S1). Because mineral N in the soil was not measured at the beginning of the experiment, the previous harvest values were used as default values at the beginning of the simulations. Since bean parameters were not available for Nsim, the model was parametrized using default parameters for pea (*Pisum sativum*).

We determined ^15^N and ^14^N levels in maize and bean tissues. The ^15^N abundance was calculated as atom% (percentage of total N) and defined as %^15^N = 100 × (^15^N)/(^15^N + ^14^N) (Marmagne et al. 2022). The ^15^N/^14^N ratio (*d15N*) was determined for the vegetative parts (*d15NV*) and propagules (*d15NP*) of maize and bean using the ^15^N abundance method (Shearer and Kohl, 1988), with the following equation:

*d15N* = 1000[(%^15^N - %^15^N_atm_)/( %^15^N_atm_)]

where %^15^N_atm_ is the percentage of ^15^N in the atmosphere, a constant value of 0.3663.

Similarly, the ^13^C/^12^C ratio (*d13C*) was calculated in the vegetative parts (*d13CV*) and propagules (*d13CP*) of maize and bean using the following equation:

*d13C* = 1000[(^13^C */*^12^C)/(^13^C */*^12^C_atm_)-1]

where ^13^C */*^12^C_atm_ is the actual ratio of ^13^C to ^12^C of the atmosphere, a constant equal to 0.011142.

Bean symbiotic nitrogen flxation was estimated from *d15N* by calculating the percent nitrogen derived from the atmosphere (*d15Ndfa*) for the vegetative parts (*d15NVdfa*) and the propagules (*d15NPdfa*) separately, using the following equation (Shearer et al. 1980):

*d15Ndfa* = 100[(*d15N_non.fix_* - *d15N_fix_*)/(*d15N_non.fix_*– B)]

where *d15N_non.fix_* is the *d15N* of a non-fixing plant such as maize, estimated from the average value of maize *d15N* between the replicates of each farm. B = *d15Nfix* when plant N only comes from atmospheric N (when plants are grown in N-free soil). Here, the B value of the *d15N* common bean was set to -1.988. This value was proposed by Farid and Navabi (2015) as the average of *d15N* measurements of 20 randomly selected bean genotypes (Peoples et al. 2002).

##### Statistical analyses

Analysis of variance (ANOVA) with a linear model including farm and cropping system as fixed effects (R package “stats”) was used to compare both farms and cropping systems (sole-cropped bean, sole-cropped maize or maize-bean intercropping) based on the 13 soil properties, simulated nitrogen mineralization (Nsim), irrigation and N fertilization (Nin).

Plant phenotypes were assessed separately for maize and bean, first using Principal Component Analysis (PCA) and ellipse plots using the factoextra R package, where ellipses represent the mean 95% confidence interval (https://github.com/kassambara/factoextra). Second, the effect of farm and cropping system on plant phenotypes were examined through ANOVA. To assess the farm-by-cropping system interaction for each crop, the analysis was restricted to farms practicing both sole-cropping and intercropping (F3, F5, F6, F7 for beans, and F1, F2, F4, F6, F7 for maize), and used the three plots per field as replicates.

For multiple testing, PCA plots were used to determine the effective number of independent traits as the minimal number of principal components required to explain 90% of the variation in soil, bean, and maize, respectively. Each *P* value was adjusted by multiplying with the corresponding number of components.

To investigate interactions between maize and bean, correlations were examined between phenotypic traits of both species using intercropping data, correcting for multiple testing through False Discovery Rate (FDR; Benjamini and Hochberg 1995) with a 20% threshold. Negative correlations are expected for competition between species, while positive correlations suggest beneficial interactions.

#### 2.1.3 Diversity analyses of OTUs from soil samples

Metabarcoding analysis of the soil bacterial community was done using 39 composite samples from the three plots in each of the 16 fields i.e., non-rhizospheric samples from the 16 fields (16 samples), plus rhizospheric samples (i) separately for maize and bean in the seven intercropping fields (14 samples) and (ii) for sole-cropping fields (9 samples). All samples were taken at 5-15 cm depth using sterile scalpels. For rhizospheric soil, 2 or 3 plants of each species were sampled per plot (6-9 plants of each species per field) and soil in contact with roots was taken from 2 to 3 different roots per plant. Composite samples for each species were made of soil from roughly 18 roots per field (∼1 g per composite sample). Likewise, non-rhizospheric soil was taken from 2 to 3 root-free soil samples in each of the 3 plots (∼1 g non-rhizosphere soil per composite sample).

Each composite sample was placed in a 2 ml ZR BashingBead Lysis tube containing beads and ZR stabilization reagent (Zymo Research, Irvine, CA) and was mixed immediately in the field using a bead-beating TerraLyzer lysis system (Zymo Research) for 30 s. DNA extraction was performed using the FastDNA kit (MP Biomedicals, Santa Ana, CA), following the manufacturer’s instructions. We sequenced 16S rRNA genes using Illumina MiSeq technology (2 × 300 bp). Sequence trimming and quality control (removal of sequences < 150 bp or with ambiguous base calls) were performed by MR DNA (Shallowater, TX). The remaining sequences were denoised, Operational Taxonomic Units (OTUs; defined at 97%) identified, and chimeras removed. Final OTUs were taxonomically classified (Datasets S5 and S6) using BLASTn against RDPII (http://rdp.cme.msu.edu). Datasets without singletons (i.e., sequences found once among the 39 samples) were used to generate rarefaction curves.

Data analysis was conducted using the phyloseq R package (McMurdie and Holmes 2013). At each taxonomic level, the 10 most abundant taxa in the entire dataset (and their proportions) were obtained using percentage normalization. For other analyses, the rarefy_even_depth function was used to normalize sample sizes based on the smallest sample size, and alpha diversity indices i.e., observed diversity (number of unique taxa) and Chao1 (taxa richness) were computed using the plot_richness function. Unifrac distances (dissimilarity between microbial communities) were calculated between samples and visualized using Non-metric Multi-Dimensional Scaling (NMDS). Permutational Multivariate Analysis of Variance (PERMANOVA) was employed to test differences using the adonis2 function from the vegan R package (https://github.com/vegandevs/vegan), with sample distances as input. The unique and common OTUs between conditions were determined on non-normalized data using Venn diagrams created with the limma package (Ritchie et al. 2015). Furthermore, variations among conditions were assessed using Between Class Analysis (BCA).

Differential analysis was done based on OTU counts using negative binomial (a.k.a. Gamma-Poisson) distribution and Wald significance tests to identify discriminant OTUs (using FDR *Q* value) between sole-cropped and intercropped conditions for maize and bean, separately, which was performed using the DESeq2 R package (https://github.com/thelovelab/DESeq2).

Subsequently, we compared OTU composition among soil types and farming practices using co-inertia analysis. The *RV* coefficient between the two datasets was computed and tested using Monte-Carlo tests with 10,000 replicates. BCA and co-inertia analyses were conducted using the ade4 R package (https://adeverse.github.io/ade4/).

### 2.2 Experimental assays of maize-bean intercropping in Saclay

#### 2.2.1 Experimental design and field management

The effect of maize-bean intercropping at the transcriptomic level was studied in a controlled field experiment in Saclay (France) in 2022. A single bean genotype (B, the Tarbais variety Alaric) was associated with maize DG0801 (M1, INRAE hybrid used as a control for low-nitrogen trials in southwestern France) and LG30.275 (M2, commercial hybrid used by INRAE as a control in northern France). The three genotypes (B, M1, M2) were grown in sole-cropping along with the two possible intercropping (M1B and M2B) in a three-block randomized design. Each block therefore contained five randomized plots with 160 cm between plots. Each plot encompassed two 5.2-m-long rows of 12 plants for sole-cropping with 24 cm intra-row spacing and 80 cm inter-row spacing, and similarly for intercropping, albeit with 12 maize and 12 beans per row. In sole-cropping plots, the density was 2.36 plants·m^-2^, while in intercropping plots, each species was sown at 2.36 plants·m^-2^, leading to doubled plant density, which is close to on-farm practice for intercropping.

Before sowing, nitrogen fertilization was performed as recommended by the Tarbais bean cooperative (residual nitrogen of 26 kg·ha^-1^ after winter and application after sowing of 75 kg·ha^-1^ of nitrogen as 27% ammonium nitrate). Such nitrogen supply corresponded to a grain yield target of 30% of their potential for commercial maize hybrids. No other chemical treatment was applied. Maize was sown on April 28^th^, 2022. Beans seeds were inoculated with a mix of symbiotic rhizobia (LEGUMEFIX Phaseolus Inoculant, Legume Technology LTD, Nottingham, UK) and sown after maize emergence on May 24^th^, 2022. Stakes were used to support bean sole-cropping. Drought occurred after bean sowing, with only 94.5 mm cumulated rainfall for the first three months, as opposed to the 188 mm average rainfall observed over the 20 previous years. Manual irrigation was applied once during the experiment.

#### 2.2.2 Comparison of gene expression between sole-cropping and intercropping

Leaf sampling was performed on the morning of July 11^th^, 2022, with a collection time minimized to ensure physiological consistency across plants. This date coincided with the flowering stage for M2, an advanced vegetative stage for M1, and an early flowering stage in beans where nodules are likely most active (Peña-Cabriales and Castellanos 1993). We pooled 1 leaf per plant for 10 randomly selected plants per plot, for a total of 21 samples (9 sole-cropping samples i.e., 3 plots × 3 blocks, and 12 intercropping samples i.e., 2 plots × 3 blocks × 2 species; Dataset S7). We sampled the 11^th^ leaf for maize, and the terminal leaflet of the last fully developed leaf for the bean. We retained 2 cm^2^ of the central portion of the blade without the midrib from the 10 leaves. Samples were flash-frozen and stored at -80 °C. Traits measured on maize and bean plants are given in Dataset S8.

Total RNAs were extracted from 100 mg of tissue with the Nucleospin RNA plant kit (Macherey-Nagel, Düren, Germany) following the manufacturer’s instructions. Extracted RNAs were checked for purity and quantity with a bioanalyzer (Agilent, Santa Clara, CA). Barcoded libraries were constructed using QuantSeq 3’RNA-Seq Library Prep Kit FWD for Illumina (Lexogen, Vienna, Austria) with Unique Molecular Identifiers (UMIs) to reduce amplification biases, and sequencing was conducted by the POPS transcriptomics platform (IPS2, Université Paris-Saclay). QuantSeq 3’RNA-seq produces a single read per transcript close to the 3′ end of polyadenylated RNAs. From 3.4 to 17.4 million 75-bp reads were obtained per sample (Dataset S7).

Pre-processing steps and read counts included deduplication of reads based on UMIs with UMI-tools v1.0.1 (Smith et al. 2017). Reads with UMI base quality score falling below 10 were removed. Removal of adapter sequences and quality trimming were performed with BBduk from the BBmap suite (v38.84, https://sourceforge.net/projects/bbmap/) with the options k=13 ktrim=r useshortkmers=t mink=5 qtrim=r trimq=10 minlength=30. The nf-core/rnaseq pipeline version 3.10.1 (Ewels et al. 2020; Patel et al. 2024, https://doi.org/10.5281/zenodo.7505987) was then used, as implemented with “nextflow” version 22.10.6 (Di Tommaso et al. 2017). Fastq files were sub-sampled with fq version 2022-02-15 (https://github.com/stjude-rust-labs/fq) and strandedness was automatically inferred using Salmon version 1.4.0 (Patro et al. 2017). Trimmed reads were mapped using STAR (v2.7.9a, Dobin et al. 2013) with default parameters, on the Zm-B73-REFERENCE-NAM-5.0 maize reference genome (https://phytozome-next.jgi.doe.gov/info/Zmays_Zm_B73_REFERENCE_NAM_5_0_55) and on the *Phaseolus vulgaris* v2.1 bean reference genome (https://phytozome-next.jgi.doe.gov/info/Pvulgaris_v2_1). Quantification was made with Salmon version 1.4.0, and the file “salmon.merged.gene_counts.tsv” was used for subsequent analysis.

From the raw read counts, reproducibility among replicates (blocks) was verified by computing R^2^. We removed library B1_M2B1_1, which was identified as an R^2^ outlier in the correlations with block 2 and block 3 (Figure S2). Standard filtering and normalization were performed using EdgeR (Lun et al. 2016). Scaling factors for each library were computed using the Trimmed Mean of M-values (TMM) normalization method (Robinson and Oshlack 2010) implemented in EdgeR (function calcNormFactors) with the following parameters: min.count=10, min.total.count=15, large.n=3, min.prop=0.6. Multi-Dimensional Scaling (MDS) on the resulting normalized counts was performed to compute pairwise distances between libraries.

For differentially expressed (DE) genes, EdgeR was further used to estimate the trended dispersion parameters (function estimateDisp), fit a quasi-likelihood negative binomial generalized log-linear model (glmQLFit), and a quasi-likelihood F-test was employed (Lun et al. 2016) (function glmQLFTest) to compute contrasts and associated *P* values for each transcript. The *P* values were adjusted to control for 5% FDR (Benjamini and Hochberg 1995). First, using only the intercropped beans, we contrasted maize genotype M1 vs M2. Second, we contrasted sole-cropping vs intercropping systems to identify DE genes and calculated the corresponding adjusted *P* values. To do so, we decomposed the expression variance into cropping system (sole-cropping vs. intercropping) and block effects for beans and into cropping system, genotype, block, and cropping system-by-genotype interaction effects for maize.

## 3. RESULTS

### 3.1 Soil properties and farming practices vary between farms

Although the survey focused on a local area around the city of Tarbes, soil properties and farming practices (including plant varieties) may vary among the 7 study farms, which could influence plant growth and plant-plant interactions. Soil properties were compared. Overall, the soils of the 16 fields were all loamy, but their contents varied from 17.1% to 30.0% for clay, 36.1% to 62.0% for silt, and 10.3% to 44.8% for sand (Figure S3). Significant differences were found among farms for 6 of the 13 soil variables (Dataset S9), and variability between fields was mainly due to soil fertility variables such as total N content (N), organic matter content (OM), C:N ratio (CN) and Cation Exchange Capacity (CEC), as exhibited on the first principal component of the PCA (Dim1, Figure 2a).

**FIGURE 2.**
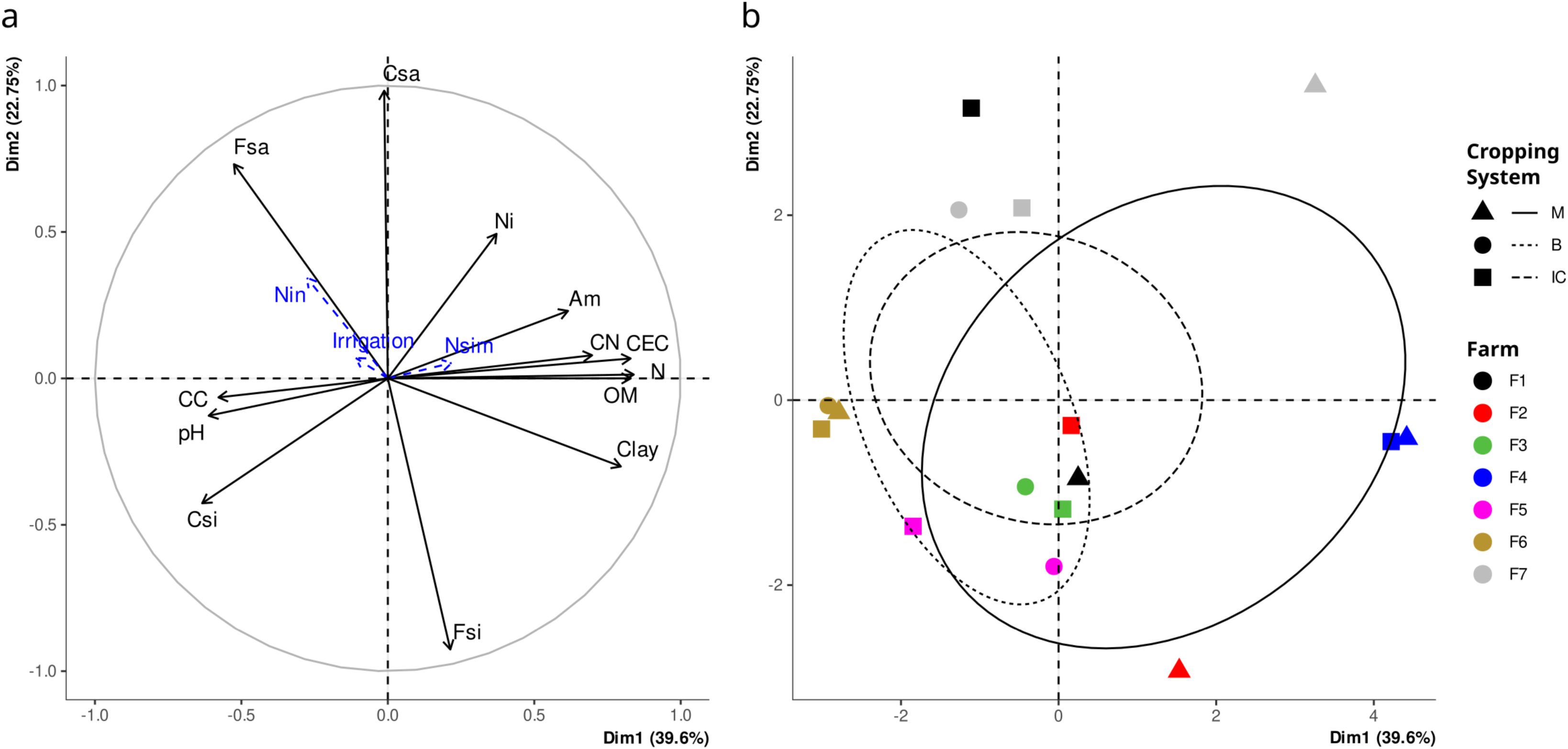
Principal Component Analysis (PCA) computed on 13 soil variables per field. The two first axes are shown. (a) Correlation circle of the 13 soil variables with three supplementary variables added to the PCA (in blue) i.e., Nsim (which estimates soil nitrogen mineralization), Irrigation, and Nin (nitrogen units supplied during the growing season). (b) Projection of fields with colors indicating farms and symbols indicating the cropping system i.e., maize sole-cropping (M), bean sole-cropping (B), or maize-bean intercropping (IC). Initial variables used in the PCA are Ni (0.5M KCl-extractable NO_3_^-^ N content), Am (0.5M KCl-extractable NH_4_^+^ N content), Clay, Fsi (Fine silt), Csi (Coarse silt), Fsa (Fine sand), Csa (Coarse sand), pH, CEC (Cation Exchange Capacity), N (Total Nitrogen), CN (Carbon:Nitrogen ratio), OM (Organic Matter) and CC (Calcium Carbonate).

Regarding farming practices, the sowing dates for the different farms ranged from April 15^th^ to June 10^th^, 2017. Since manual harvest of beans requires vast space between rows, sole-cropped maize was cultivated at a much higher density (8 plants·m^-2^) compared with intercropped maize and bean (2.5 plants·m^-2^ for each) and sole-cropped bean (2.5 plants·m^-2^). Except for the organic farm F4, with low N-fertilization (50 kg N·ha^-1^ for sole-cropped maize and no fertilization for intercropped maize and bean), N-fertilizer input was on average 224 N units·ha^-1^ for sole-cropped maize, 130 N units·ha^-1^ for sole-cropped bean, and 135 N units·ha^-1^ for intercropped maize and bean. Farm F4 did not practice irrigation. For the other farms, irrigation along the growing season was, on average, 181 mm for sole-cropped maize, 150 mm for sole-cropped bean, and 117 mm for intercropped maize and bean.

Among the three variables related to farming practices (Nsim, Nin, and irrigation), the simulated soil nitrogen mineralization from organic matter (Nsim) and the N fertilization (Nin) differed among farms (Dataset S9). Notably, the only organic farm in our panel (F4) stood out as an outlier along the first principal component of the PCA, suggesting superior soil fertility as expected from the longer crop rotation and high pre-sowing manure inputs in this farming system (Figure 2b). Conversely, farm F6 (which used the highest quantity of nitrogen fertilizers; Dataset S1) displayed low values for fertility soil variables (Figure 2). Nsim was positively correlated with N, OM, and CEC (*r* > 0.63), while it was negatively correlated with irrigation (*r* = -0.79, *P* = 0.0003, Dataset S10). Additionally, Nsim exhibited a negative correlation with N fertilization (*r* = -0.27, *P* = 0.037), as expected, indicating that soils with greater soil mineralization require less fertilization.

### 3.2 Agronomic practices and cropping system shape soil bacterial communities

Beyond the well-known bean–rhizobia symbiosis, root interactions with rhizospheric microbiota influence plant development, prompting comparison of rhizospheric and non-rhizospheric bacterial communities under sole- and intercropping systems. A total of 4,632,917 16S rRNA sequences were obtained. Their abundance varied from 80,394 to 160,393 sequences (Dataset S5). Sequence assignation resulted in 12,171 bacterial OTUs, classified at seven taxonomic levels (Dataset S6). When combining data from all farms, the most prevalent phyla were *Proteobacteria*, *Actinobacteria*, *Bacteroidetes*, *Chloroflexi*, and *Firmicutes* (Figure S4).

Interestingly, we observed no significant difference in alpha diversity between maize and beans. However, intercropping conditions displayed a trend toward higher alpha diversity (Observed OTUs and Chao1 indexes) irrespective of the rhizosphere/non-rhizosphere status for both maize and bean soils (Figure S5). This increase in diversity under intercropping conditions was significant for soils associated with maize only (FDR *Q* = 0.014). The observed diversity of non-rhizosphere soil in intercropping conditions appeared to be intermediate between the intercropped maize and intercropped bean non-rhizosphere soils.

We examined distances between samples considering all OTUs (Figure S6). We revealed closer proximity among samples originating from the same farm (“farm” effect at *P* < 0.001 in PERMANOVA, Dataset S11), regardless of soil conditions (rhizosphere vs non-rhizosphere), plant species (maize vs bean) or cropping system (intercropping vs sole-cropping). OTUs whose counts differed among samples separated rhizospheric from non-rhizospheric samples with a more precise pattern for beans than for maize (Figure 3). The second axis of the BCA differentiated samples by cropping system for both crops (Figure 3).

**FIGURE 3.**
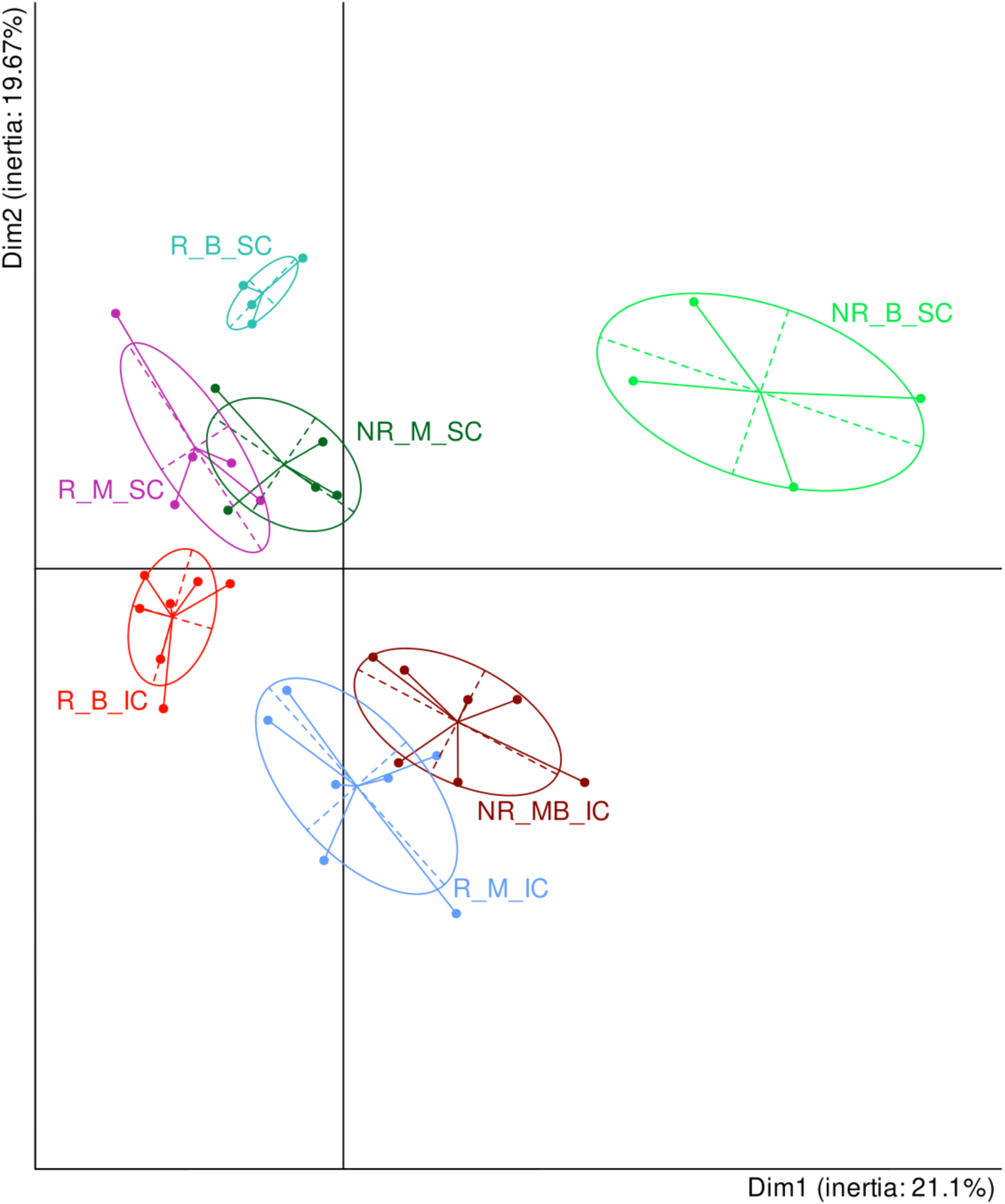
Between Class Analysis (BCA) computed on 11,586 OTUs. Samples are named according to rhizosphere (R) or non-rhizosphere (NR) soil, followed by maize (M) or bean (B), and then sole-cropping (SC) or intercropping (IC). Projections on the first two dimensions are presented, with the corresponding percentages of total inertia explained by these axes.

Co-inertia analysis unveiled a significant association between OTU composition and the 13 soil variables, irrigation, N input, and Nsim (correlation coefficient *RV* = 0.58; *P* = 0.025, Figure S7). The first dimension of soil PCA (Figure 2) exhibited a negative contribution to the first dimension of co-inertia (-0.93), while the second dimension of soil PCA contributed positively to the second dimension of co-inertia (0.93). The first two axes of the co-inertia accounted for 61.2% of the total inertia (Figure S7). Notably, Irrigation and Nsim presented the highest values of canonical weights (respectively 0.45 and -0.43, Dataset S12) that defined the second axis of co-inertia, indicating that these two variables highly impacted OTU composition. F4 and F6 emerged as the most distinct farms on the co-inertia plot along the first axis (Figure S7). Interestingly, F4 was the only organic farm in our sample, applying the lowest level of mineral inputs (Dataset S1) while displaying superior soil fertility (Figure 2).

Finally, Wald multiple comparison tests carried out with individual OTUs indicated that *Vibrio*, *Acinetobacter*, *Pseudonocardia alaniniphila*, *Hydrogenophaga palleronii* and *Pseudoxanthomona koreensis* were more abundant in sole-cropped than intercropped bean rhizosphere, while *Janthinobacterium lividum* was more abundant in the intercropped bean rhizosphere (Figure S8). In the maize rhizosphere, *Actinospica*, *Sphingomonas mali, Ktedonobacter robiniae* (previously bacterium SOSP1-30), and *Alcaligenes* were more abundant in sole-cropped samples, while *Thermoactinomyces* and *Bdellovibrio* were more abundant in intercropped samples (Figure S9). These last results need, however, to be taken with caution as the data were scarce and significance was, in some instances, driven by a single data point.

### 3.3 Cropping system impacts maize agronomic traits

The co-cultivation of beans within maize fields may affect maize development and yield in negative (through competition) or positive (through facilitation) ways, depending on plant varieties, soil properties, or farming practices. Thus, we examined the effect of the cropping system on maize agronomic traits using a multivariate analysis. We found that the first principal component explained 40.9% of the variation and was strongly associated with yield-related variables, including propagule dimensions and number (Dim 1, Figure 4a). This principal component correlated positively with nitrogen fertilization (Nin, *r* = 0.77, *P* < 0.01) and irrigation (*r* = 0.68, *P* < 0.01) and negatively with simulated nitrogen mineralization (Nsim, *r* = -0.78, *P* < 0.01). In addition, different fields of a given farm tended to cluster along the first principal component, suggesting that farming practices and local soil properties primarily determined maize yield (Figure 4b). The ellipse plot further revealed a clear distinction between sole-cropped (M) and intercropped maize (IC) along this axis (Figure 4b).

**FIGURE 4.**
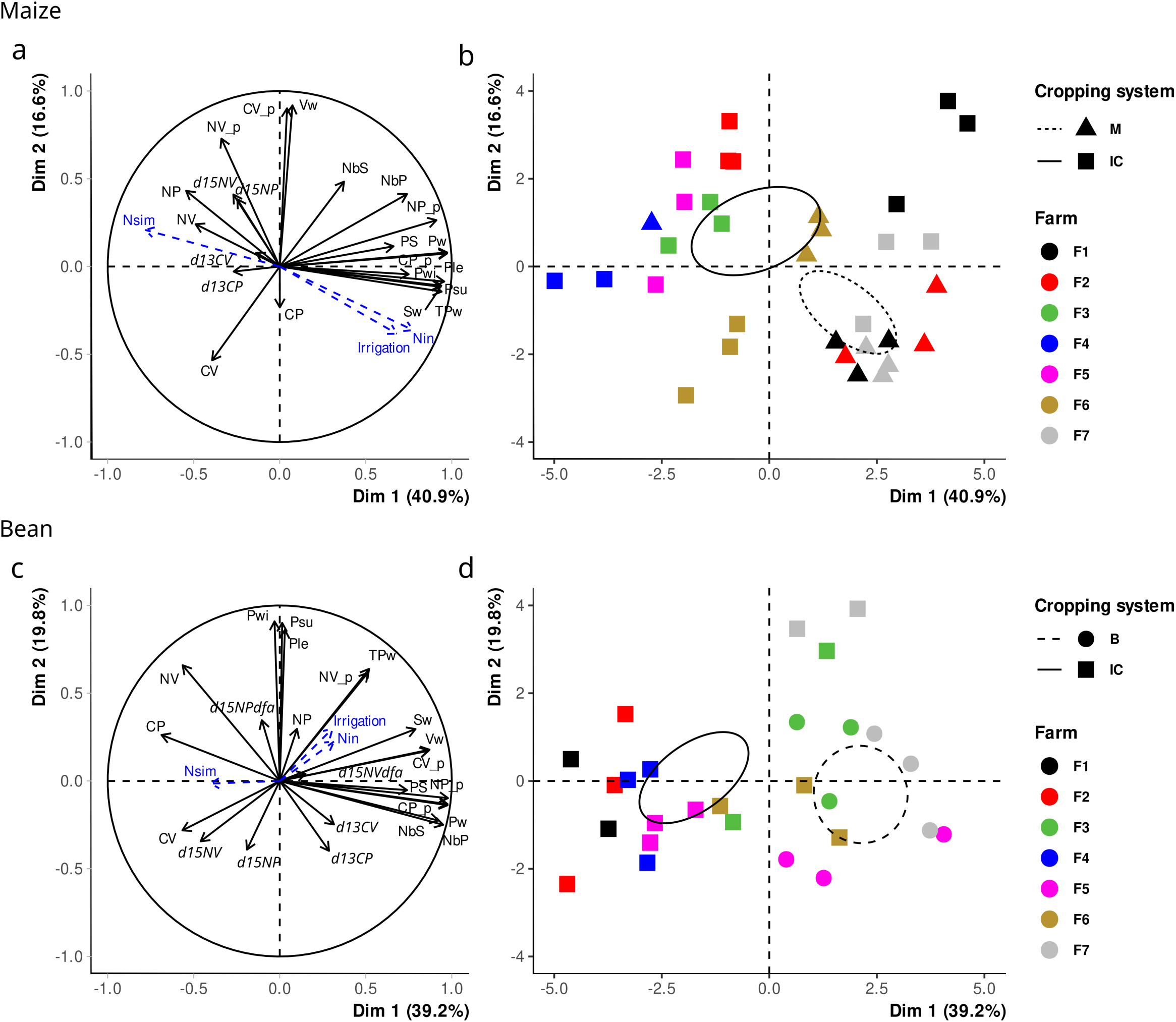
Principal Component Analysis (PCA) on maize and bean phenotypic and nutrition variables. The first two principal components (PCs) computed from 22 maize variables (Table 1) are shown with corresponding correlation circle (a) and data projection (b). Similarly, for beans, the first two PCs computed on 24 variables (Table 1) are shown with corresponding correlation circle (c) and data projection (d). The three variables, Irrigation, Nin (N input), and Nsim (simulated soil mineralization), shown in blue (a and c), were added to the PCA. In b and d, colors indicate farms, and symbols indicate cropping system with corresponding ellipses i.e., sole-cropped maize in b (M, dashed lines), sole-cropped bean in d (B, dashed line), intercropped maize in b and intercropped bean in d (IC, full lines).

The difference between cropping systems was stable across farms for three traits, intercropping being associated with higher values of two N-nutrition-related traits (NP and *d15NV*) and lower values of kernel weight per ear (Sw), regardless of the farm (Figure 5, Dataset S13). Although showing significant farm-by-cropping system interaction, the percentage of carbon in vegetative parts (CV) exhibited consistently smaller intercropping values than in sole-cropping (Figure S10). For eight other traits, the difference between cropping systems varied among farms (i.e., significant farm-by-cropping system interactions, Dataset S13).

**FIGURE 5.**
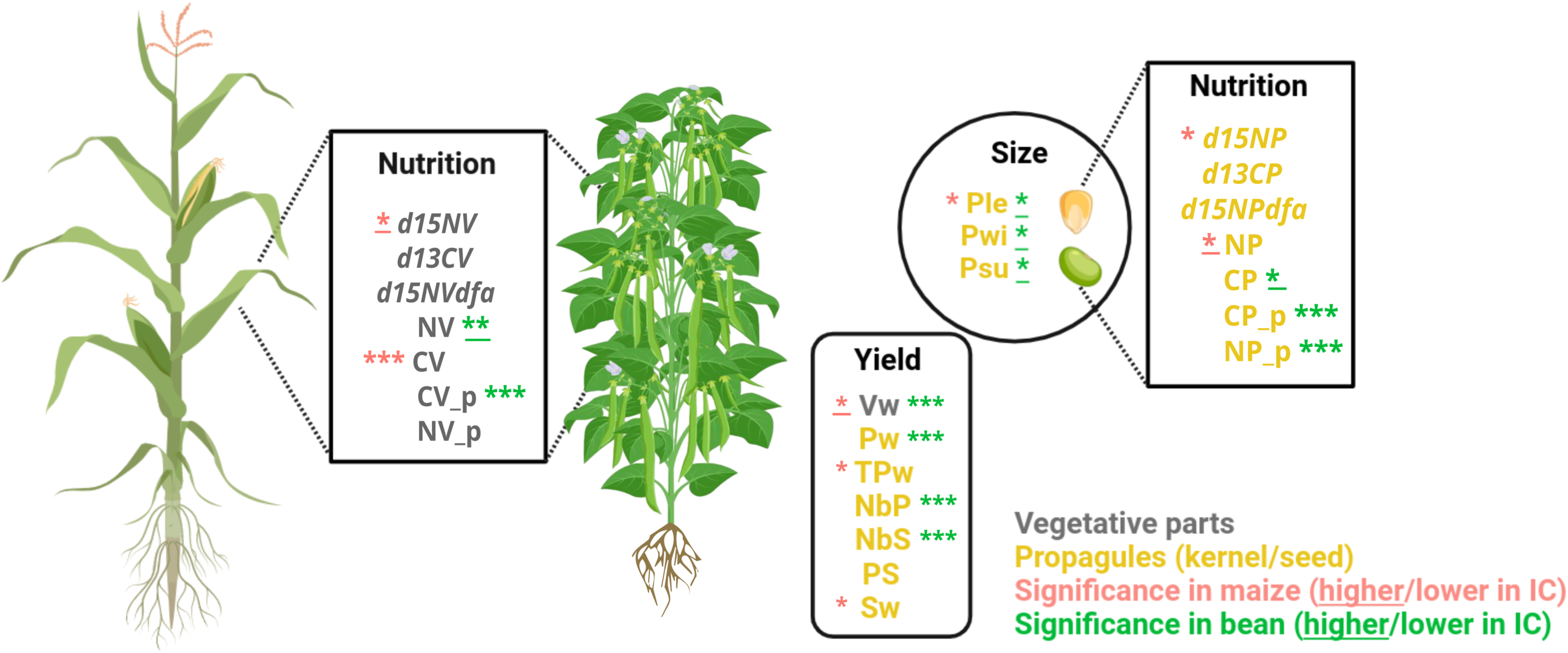
Effect of the cropping system on maize and bean phenotypic and nutrition variables. We tested the effect of the farm and the Cropping system (sole-cropping versus intercropping) on phenotypic and nutrition variables using a two-way analysis of variance model with interaction. Variables impacted by intercropping are *d15NV* (Delta ^15^N among total N of the Vegetative parts), NV (Percent Nitrogen in Vegetative parts), CV (Percent Carbon in Vegetative parts), CV_p (Per plant Carbon content in Vegetative parts), Vw (Vegetative parts dry weight per plant), Pw (Propagule dry weight per plant), TPw (Thousand Propagules dry weight), NbP (Number of Propagules produced per plant), NbS (Number of Sets of propagules per plant), Sw (Average propagule weight per Set), Ple (Average Propagule length), Pwi (Average Propagule width), Psu (Average Propagule surface), NP (Percent Nitrogen in Propagules), CP (Percent Carbon in Propagules), CP_p (Per plant Carbon content in propagules) and, NP_p (Per plant Nitrogen content in propagules). Pink asterisks indicate the significance of tests performed on maize traits, and green asterisks those for bean traits. Asterisks are underlined when the value is higher in the intercropping than in the sole-cropping system. Significant *P* values for the cropping system effect are indicated by *, **, and *** for *P* < 0.05, *P* < 0.01, and *P* < 0.001, respectively.

### 3.4 Cropping system impacts bean agronomic traits

Sole-cropping of climbing bean varieties increases costs for the farmer, who must install expensive, single-use nets in the field. Here, we did not consider these aspects but focused on the effect of maize presence/absence on bean agronomic traits. First, multivariate analysis of the cropping system effect on bean agronomic traits indicated that the first two principal components explained over 59% of the variation (Figure 4c). The first principal component was associated with yield and nutrition traits (Pw, NbS, NbP, PS, CP_p, CV_p, NP_p, Vw), while the second one was linked to propagule size (Pwi, Ple and Psu). Sole-cropped (B) and intercropped (IC) beans were separated along the first principal component (Figure 4d). Second, intercropping negatively affected yield components across all farms, although the intensity varied, as indicated by significant farm-by-cropping system interactions (Figure S11). This included vegetative weight (Vw), seed weight (Pw), the number of seeds per plant (NbP), and the number of pods per plant (NbS). A similar pattern was observed for traits related to nitrogen and carbon content per plant, such as CV_p, CP_p, and NP_p (Figure 5, Dataset S13). Interestingly, as compared to sole-cropping, intercropped beans displayed higher values for seed size traits (Ple, Pwi, Psu) as well as for nutrition-related traits (NV, CP) across all farms (Figure 5, Dataset S13).

### 3.5 Competition prevails in maize-bean intercropping

Correlations between maize and bean traits were computed to explore the interplay between the crops, with positive correlations expected under facilitation and negative correlations under competition. Of the 528 correlations (22 traits for maize and 24 traits for bean), 47 were significant, among which 34 were negative and 13 were positive (Figure S12), pointing toward competition rather than beneficial interactions. Considering the subset of traits impacted either in the maize or the bean by intercropping (Figure 5), we found that maize traits related to nitrogen nutrition (*d15NV*, *d15NP*, CV_p ; Table 1) and vegetative dry weight (Vw) were negatively correlated both with yield-related traits (NbS, Pw, Sw, Tpw, NbP) and two nutrition-related traits (CP_p, NP_p) measured in bean (Table 1). These results indicate inter-species competition for resources such as light, water, and mineral N needed for nitrogen assimilation and CO_2_ fixation through photosynthesis, reducing bean yield. Conversely, bean’s nitrogen percentage in vegetative parts (NV) was positively correlated with four maize traits (Pw, NbP, CP_p, *d13CP*; Table 1). This most likely resulted from an indirect effect linked to the lower seed production— less nitrogen mobilized into the reproductive organs—in intercropped beans.

### 3.6 Species competition affects bean transcriptome in controlled field assay

To further assess the impact of intercropping on plant growth and gene expression, we conducted a controlled field assay at the INRAE plant breeding station of Saclay (northern France). Agronomic data revealed that 6 of 13 yield and development traits exhibited lower values in the intercropped bean as compared to sole-cropped bean, i.e. total seed mass per plant (reduced by 87%; Figure 6a), number of seeds per plant (reduced by 89%), number of pods per plant (reduced by 89%), width of the primary leaflets (reduced by 19%), foliage density (reduced by 56%), and plant height (reduced by 59%; Figure 6b) (Figure S13). Although the two maize varieties strongly differed in all these traits (except for the number of ears per plant and the number of lodged plants), both were unaffected by the cropping condition (Figure 6c & 6d and Figure S14).

**FIGURE 6.**
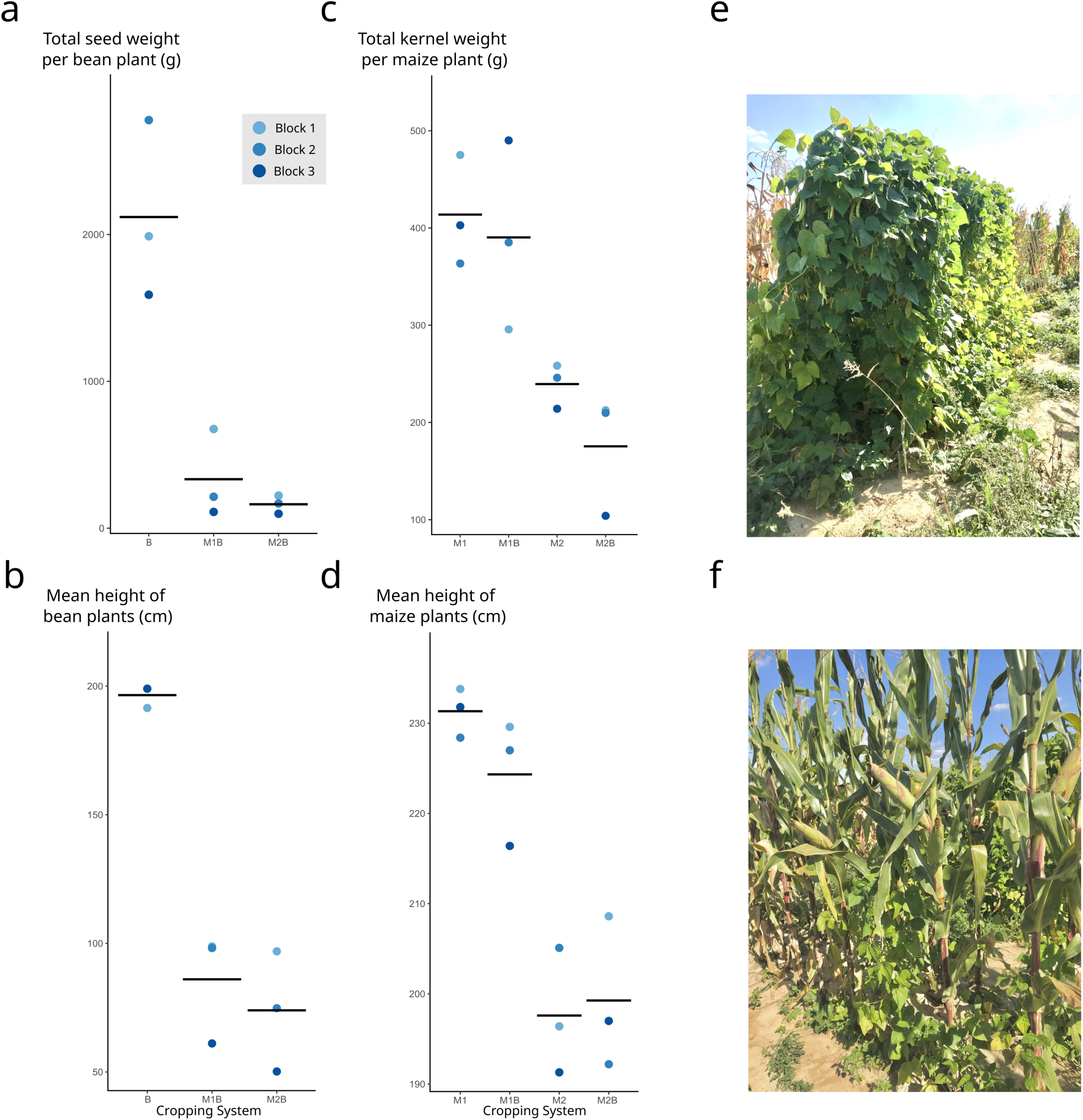
Development and yield of maize and bean plants under sole-cropping and intercropping in the field assay at Saclay. Stripcharts display the total seed weight per bean plant (a) and the height of bean plants (b), the total kernel weight per maize plant (c), and the height of maize plants (d). Blue dots represent the average value of the three blocks, and horizontal lines indicate the means. Data are shown in a and b for sole-cropped maize genotype 1 (M1), sole-cropped maize genotype 2 (M2), maize-bean intercropping (M1B and M2B), and in c and d for sole-cropped bean (B), and maize-bean intercropping (M1B and M2B). Pictures illustrate sole-cropped beans (e) and intercropped beans and maize (f).

Transcriptomic changes in sole-cropping versus intercropping stands were investigated. First, the comparison of maize gene expression between genotypes M1 and M2 showed clear distinctions between the two genotypes on the first axis of the MDS plot, which accounted for 66% of the total variation (Figure 7a). Second, differentially expressed (DE) genes were investigated, with the prediction that their number would be higher in beans than in maize since bean phenotypes (but not maize phenotypes) were severely affected by the cropping condition. Following our prediction, we did not detect any DE gene for sole-cropped vs intercropped maize. In bean, we identified only 11 DE genes between M1B and M2B and concluded on the near absence of impact of the maize variety on bean transcriptome. Thus, we considered M1B and M2B biological replicates for detecting DE genes. Of 14,993 bean genes, nearly a third (5,070 genes) were DE between cropping conditions (first MDS axis, Figure 7b).

**FIGURE 7.**
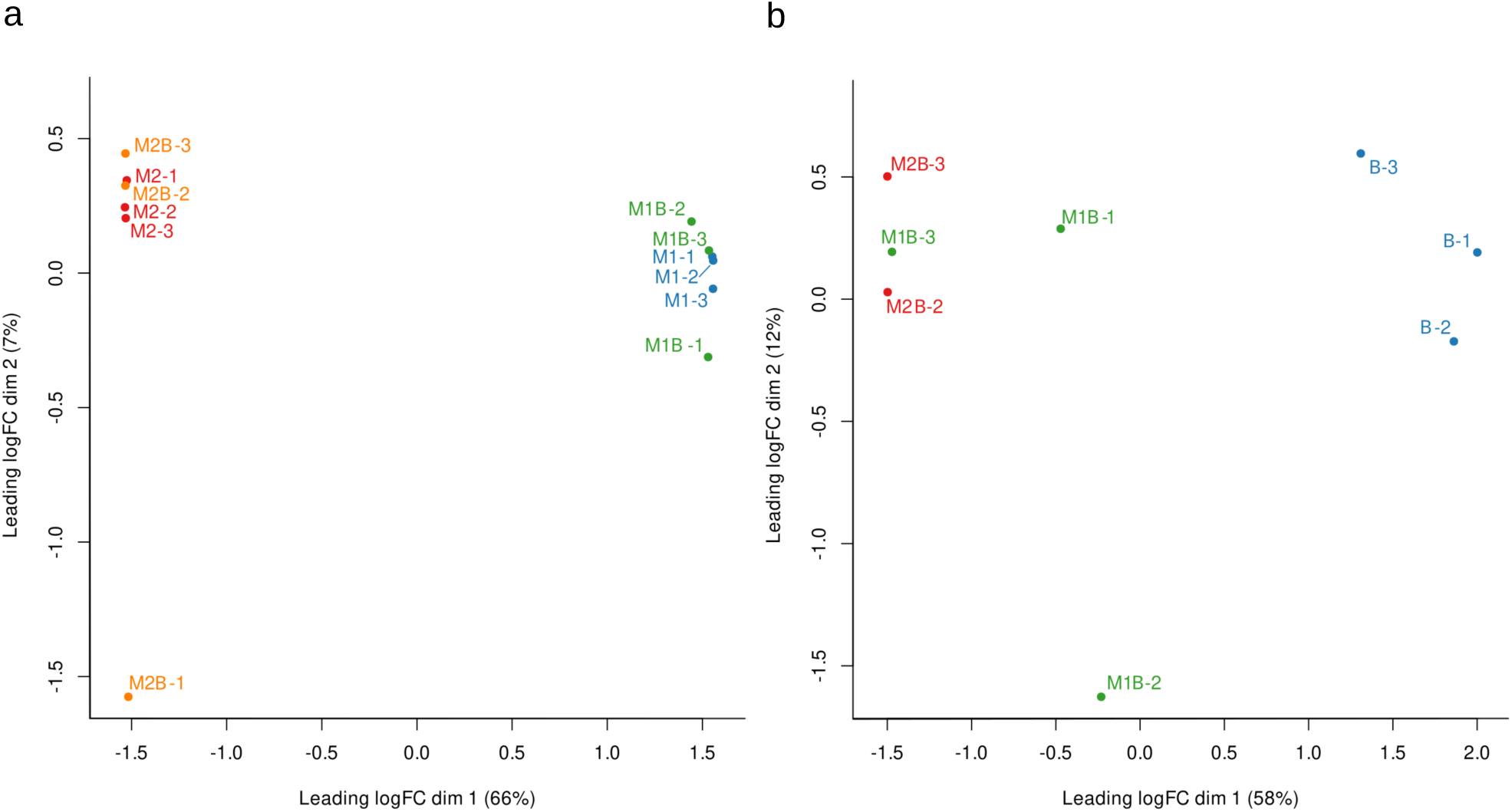
Multi-Dimensional Scaling (MDS) plots of distances between gene expression profiles in sole-cropped and intercropped maize and bean. The distances observed between samples in the two MDS plots approximate the typical log_2_ fold changes between the samples. MDS plots were computed from 16,021 genes for maize samples (a) and 14,993 genes for bean samples (b). Colors and labels indicate the cropping system and the sample for sole-cropped maize of the two different genotypes (M1 and M2), sole-cropped bean (B), and maize-bean intercropping (M1B and M2B). The last digit in the labels indicates the corresponding block (1, 2, or 3). The percentage of variation explained by each dimension is shown.

## 4. DISCUSSION

Intercropping systems are gaining popularity, and they are appreciated for their low-input requirements and yield in low-input settings. Such positive effects result from beneficial plant-plant interactions (Fréville et al. 2022). In this study, we undertook a multidisciplinary approach to get insights into maize-bean intercropping, a practice that was recently revived in southwestern France for the cultivation of the Tarbais bean. Our study’s primary strength, yet limitation, lies in the direct sampling of plants from on-farm fields. This approach introduces numerous confounding factors, such as maize variety, inputs, and abiotic and biotic environmental factors, which are challenging to control. Here, we captured these factors collectively as the “farm” effect.

Soil analyses revealed notable differences in fertility and pH across farms (Dataset S9), while soil texture remained relatively consistent (Figure S3). Globally, we observed a positive correlation between nitrogen mineralization by microorganisms and soil fertility, i.e., with total N content (*r* = 0.64; *P* < 0.01) and organic matter content (*r* = 0.64; *P* < 0.01), alongside a negative trend with nitrogen fertilizer application. These correlations driven by the organic farm suggest that farmers compensate for poor soil fertility with nitrogen inputs, and they also highlight the intricate connections between farming practices, soil fertility, and the capacity of microorganisms to release nutrients into the soil.

While it is well known that bulk bacterial communities are shaped by various soil variables— including texture (Girvan et al. 2003), nitrogen content (Zeng 2016), pH (Lauber et al. 2008), and water (Li 2021), the rhizosphere communities are also influenced by crop genotypes (Bouffaud et al. 2016) and physiology, which collectively determine the compounds released near roots as exudates (Ulbrich et al. 2022). Due to the complexity of interactions, studying the impact of intercropping on bacterial composition remains challenging (Duan et al. 2024). Here, we found a trend towards higher diversity in soil samples obtained from intercropping than sole-cropping (significant in maize; Figure S5), a pattern observed in sugarcane–soybean intercropping (Malviya et al. 2021). Interestingly, alpha diversity levels of maize and bean soils were not different, meaning that the greater diversity observed in intercropped maize more likely results from a complementarity effect rather than a selective effect (Loreau and Hector 2001). The combination of root exudates released by both crops may enhance the diversity of the associated bacterial communities. In addition, we found in bean that two of the OTUs that discriminated the cropping systems include plant growth-promoting rhizobacteria (Figure S8). The *Acinetobacter* spp., which were predominant in sole-cropped bean rhizosphere, comprise strains that can produce indole-3-acetic acid (IAA), siderophores, gibberellin, antibiotics, biosurfactants/bioemulsifiers, and solubilize phosphate, potassium, and zinc (Rokhbakhsh-Zamin et al. 2011). Also, *Janthinobacterium lividum* was more abundant in the bean rhizosphere when intercropped with maize. Interestingly, strains of this species can inhibit several soil-borne pathogens, including *Pythium ultimum*, *Rhizoctonia solani*, and *Rhizoctonia oryzae*, and display biocontrol activity in the wheat rhizosphere (Yin et al. 2021).

In cereal-legume intercropping, cereal roots develop more quickly, utilizing nitrogen before legumes, stimulating nodule formation and nitrogen fixation in beans (Hauggaard-Nielsen et al. 2008). Although we did not conduct a detailed examination of root nodules, initial observations revealed no nodules on the bean in topsoil. This is likely caused by the sampling timing (close to harvest), which is not ideal for making such observations but suggests limited symbiotic nitrogen fixation despite the nodulation capability of Tarbais beans (Laguerre et al. 2006). This could be due to two factors: the non-restrictive nitrogen fertilization conditions, which can inhibit nodulation (Voisin et al. 2002), or the absence of the specific rhizobial strains needed for nodulation in the soil (Yan et al. 2014). Because we evidenced the presence of *Rhizobium leguminosarum* (Dataset S6), involved in bean nodulation, our findings suggest that years of conventional practices of maize monoculture without a Fabaceae cover crop may have diminished the potential benefits of plant-microbial interactions in these fields.

To assess the potential beneficial interactions between maize and beans in intercropping, we examined the effect of intercropping on plant nutrition and development. We did not compute the Land Equivalent Ratio, often used to compare production in intercropping versus sole-cropping (Mead and Willey 1980), because the farming practices differed between cropping conditions. Pairwise correlations between traits of the two crops revealed that most significant correlations were negative, indicating that competition outweighed beneficial interactions. Notably, yield components in both crops were negatively affected by intercropping, with beans experiencing greater reductions compared to sole-cropping situations (Figures S10, S11, and S12). Interestingly, however, size traits of bean seeds (length, width, and surface) were larger in intercropped than sole-cropping, indicating that competition with maize translates into beans producing fewer pods and seeds but larger seeds. This pattern illustrates the trade-off in resource allocation between fecundity and seed size, the latter providing offspring with more resources (Aarssenand and Keog 2002) and, therefore, a potential competitive advantage during early development, as shown in other species (Houssard and Escarré 1991). In intercropping, maize also displayed increased nitrogen acquisition in kernels and higher ^15^N abundance in vegetative tissues (Figure 5), suggesting an enhanced capacity of maize to mobilize soil nitrogen – the elevated ^15^N levels indicate that maize primarily relies on soil nitrogen. This contrasts with beans, which showed a higher nitrogen percentage in vegetative tissues (Figure S12). The reduced capacity of certain beans to compete with maize for soil nitrogen (Dawo et al. 2009) likely explains their lower yield. In this setting, nitrogen fixation was limited, as evidenced by no significant increase in nitrogen fixation in intercropping. Hence, enhanced nitrogen in vegetative parts of beans may reflect a reduced demand for nitrogen in seeds due to less seed production and a lower dilution of nitrogen assimilates. These findings suggest that competition between species dominates the observed resource allocation patterns, with maize benefiting more from shared soil nitrogen resources while beans face limitations.

Finally, the field assay in Saclay provided a first glimpse into the characterization of the molecular signatures of maize-bean intercropping (Becker et al. 2023). Our results show that competition in intercropping was much stronger than what we observed on-farm. It is interesting to note that under such increased competition, the burden falls on the bean and results in a substantial decline in yield (∼87%) and vegetative development compared to sole-cropping beans, even though early growth of the bean remained unaffected. In contrast, maize showed no difference between cropping conditions across all measured traits. Interestingly, gene expression analysis concurred with phenotypic data, revealing no differentially expressed genes in maize but 5070 DE genes in beans. This indicated a complete rewiring of bean gene expression when suffering drastic competition.

## 5. CONCLUSION

Despite the intrinsic limitations of field surveys in farmers’ settings, including confounding effects and limited sampling, our results demonstrate that intercropping leads to plant competition that significantly impacts the agronomic traits of maize and beans in the Tarbes region. However, this impact is highly dependent on farming practices. Key factors such as crop variety, plant row arrangement and density, soil properties, and farm inputs that affect soil fertility and microbial composition play critical roles in shaping these outcomes. Differences observed between on-farm surveys and the field experiment, for instance, illustrate the potential importance of maize genotype and environmental conditions, such as soil composition and drought, in driving plant competition dynamics. These findings underscore the pivotal role of farming practices and soil fertility in mediating species interactions, extending even to the diversity of bacterial soil communities. Interestingly, intercropping tends to foster more diverse microbial communities than sole-cropping, potentially benefiting soil health. Despite this advantage, intercropping remains characterized by intense interspecific competition, particularly to the detriment of beans. This competition negatively affects both gene expression and yield-related traits in bean. However, a notable outcome is the production of larger seeds by intercropped beans compared with those grown in sole-cropping. This attribute could be valued by farmers as larger seed size may be a criterion for premium labels, guaranteeing higher market prices and stable demand.

This interdisciplinary study paves the way for recommendations on maize-bean intercropping and outlines several research perspectives. First, the genetic basis of beans in the Tarbes region is remarkably narrow, and the two bean varieties have been selected without explicit consideration of their symbiotic N-fixation potential in soils of this area, where rhizobial inoculation is not customary. Second, beans are the primary focus in this system, and selecting an appropriate maize variety is crucial. The ideal maize partner should be robust enough to support the beans but not overly competitive. Hybrids optimized for sole-cropping and mineral nitrogen absorption are unsuitable in this context. Molecular studies, such as those conducted here but expanded to other growth stages, plant organs, and varieties, could help identify the metabolic pathways and guide partner selection using genetic resources. Third, the study encourages a reevaluation of the criteria for the success of this intercropping system. Beyond productivity, traits such as the nutritional quality of seeds and/or their specific characteristics could be shaped by consumer preferences while acknowledging the positive impacts of intercropping on the agroecosystem.

## ACKNOWLEDGEMENTS

We are incredibly grateful to Hélène Corti, a retired technician who helped us collect the data, and Marie-Hélène Jeuffroy and Etienne Delannoy for insightful discussion at the start of the project. This work would not have been possible without the contribution of farmers and the Coopérative du Haricot Tarbais. Three anonymous reviewers greatly contributed to improving the manuscript. We thank Andrew Crawford, from the Academic writing center of Paris-Saclay University, for his careful proofreading of the final version of the manuscript.

## AUTHOR CONTRIBUTIONS

Conceptualization, M.C., T.F., J.E., Y.M-L., Do.M., M.I.T.; Methodology, N.V.-B., M.C., C.B., T.F., J.E., V.P., E.L.C., Da.M., Y.M.-L., Do.M., M.I.T.; Software, N.V-B, T.T., C.P.-L., T.F., Da.M., Y.M.-L., Do.M., M.I.T.; Validation, N.V.-B., L.M., T.T.; Formal analysis, N.V.-B., L.M., T.T., V.P.; Investigation, N.V.-B., B.L., C.P., M.C., M.L.G., J.C., A.M., Y.M.-L., Do.M., M.I.T.; Resources, C.P., M.C., C.B., J.C., C.P.-L., T.F., M.L.G, V.P., A.M., Y.M.-L.; Writing – Original Draft, N.V.-B., L.M., Do.M., M.I.T.; Writing –Review & Editing, N.V.-B., L.M., T.F., J.E., E.L.C., V.P., Da.M., Y.M.-L., Do.M. M.I.T.; Vizualization, N.V.-B., L.M., T.T.; Supervision, Y.M.-L., Do.M. M.I.T.; Project administration, Y.M.-L., M.I.T.; Funding Acquisition, N.V.-B., C.P.-L., T.F., Y.M.-L., Do.M., M.I.T.

## DATA AVAILABILITY STATEMENT

Raw reads from metabarcoding data have been deposited in the DDBJ/EMBL/GeneBank under accession number PRJNA 1112577 and similarly for raw reads from QuantSeq 3’RNA-seq under accession number PRJNA 1115135. Raw phenotypic data and input files to run the code have been deposited on recherche.data.gouv.fr with the DOI https://doi.org/10.57745/BZMPT9. RNAseq and phenotypic data analyses codes are available at recherche.data.gouv.fr with the DOI https://doi.org/10.57745/BZMPT9.

## SUPPORTING INFORMATION

This paper contains a supporting information file with 14 supplementary figures and 13 supplementary datasets.

## REFERENCES

Aarssen, L. W., & Keog, T. (2002). Conundrums of competitive ability in plants: what to measure? Oikos, 96(3), 531–542. 10.1034/j.1600-0706.2002.960314.x

Becker, C., Berthomé, R., Delavault, P., Flutre, T., Fréville, H., Gibot-Leclerc, S., Le Corre, V., Morel, J-B., Moutier, N., Munos, S., Richard-Molard., C., Westwood, J., Courty, P-E., de Saint Germain, A., Louarn, G., & Roux, F. (2023). The ecologically relevant genetics of plant-plant interactions. Trends in Plant Science, 28(1), 31–42. 10.1016/j.tplants.2022.08.014

Bedoussac, L., Journet, E.-P., Hauggaard-Nielsen, H., Naudin, C., Corre-Hellou, G., Jensen, E. S., Prieur, L., & Justes, E. (2015). Ecological principles underlying the increase of productivity achieved by cereal-grain legume intercrops in organic farming. A review. Agronomy for Sustainable Development, 35(3), 911–935. 10.1007/s13593-014-0277-7

Beillouin, D., Ben-Ari, T., Malézieux, E., Seufert, V., & Makowski, D. (2021). Positive but variable effects of crop diversification on biodiversity and ecosystem services. Global Change Biology, 27(19), 4697–4710. 10.1111/gcb.15747

Benjamini, Y., & Hochberg, Y. (1995). Controlling the false discovery rate : A practical and powerful approach to multiple testing. Journal of the Royal Statistical Society: Series B (Methodological), 57(1), 289–300. 10.1111/j.2517-6161.1995.tb02031.x

Bitocchi, E., Bellucci, E., Giardini, A., Rau, D., Rodriguez, M., Biagetti, E., Santilocchi, R., Zeuli, P. S., Gioia, T., Logozzo, G., Attene, G., Nanni, L., & Papa, R. (2013). Molecular analysis of the parallel domestication of the common bean (*Phaseolus vulgaris*) in Mesoamerica and the Andes. New Phytologist, 197(1), 300–313. 10.1111/j.1469-8137.2012.04377.x

Bonnain-Dulon, R., & Brochot, A. (2004). De l’authenticité des produits alimentaires. Ruralia, 14, 969. http://journals.openedition.org/ruralia/969

Bouffaud, M.-L., Renoud, S., Moënne-Loccoz, Y., & Muller, D. (2016). Is plant evolutionary history impacting recruitment of diazotrophs and *nifH* expression in the rhizosphere? Scientific Reports, 6(1), 21690. 10.1038/srep21690

Brooker, R. W., Bennett, A. E., Cong, W. F., Daniell, T. J., George, T. S., Hallett, P. D., Hawes, C., Iannetta, P. P. M., Jones, H. G., Karley, A. J., Li, L., McKenzie, B. M., Pakeman, R. J., Paterson, E., Schob, C., Shen, J. B., Squire, G., Watson, C. A., Zhang, C. C., … White, P. J. (2015). Improving intercropping : A synthesis of research in agronomy, plant physiology and ecology. New Phytologist, 206(1), 107–117. 10.1111/nph.13132

Clivot, H., Mouny, J.-C., Duparque, A., Dinh, J.-L., Denoroy, P., Houot, S., Vertès, F., Trochard, R., Bouthier, A., Sagot, S., & Mary, B. (2019). Modeling soil organic carbon evolution in long-term arable experiments with AMG model. Environmental Modelling & Software, 118, 99–113. 10.1016/j.envsoft.2019.04.004

Dawo, M. I., Wilkinson, J. M., & Pilbeam, D. J. (2009). Interactions between plants in intercropped maize and common bean. Journal of the Science of Food and Agriculture, 89(1), 41–48. 10.1002/jsfa.3408

Di Tommaso, P., Chatzou, M., Floden, E. W., Barja, P. P., Palumbo, E., & Notredame, C. (2017). Nextflow enables reproducible computational workflows. Nature Biotechnology, 35(4), 316–319. 10.1038/nbt.3820

Dobin, A., Davis, C. A., Schlesinger, F., Drenkow, J., Zaleski, C., Jha, S., Batut, P., Chaisson, M., & Gingeras, T. R. (2013). STAR : Ultrafast universal RNA-seq aligner. *Bioinformatics* (Oxford, England), 29(1), 15–21. 10.1093/bioinformatics/bts635

Duan, Y., Lei, X., Cao, Y., Liu, L., Zou, Z., Ma, Y., Zhu, X., & Fang W. (2024). Leguminous green manure intercropping changes the soil microbial community and increases soil nutrients and key quality components of tea leaves. Horticultural Research, 11(3): uhae0188. 10.1093/hr/uhae018

Ewels, P. A., Pelzer, A., Fillinger, S., Patel, H., Alneberg, J., Wilm, A., Garcia, M. U., Di Tommaso, P., & Nahsen, S. (2020). The nf-core framework for community-curated bioinformatics pipelines. Nature Biotechnology, 38(3), 271–271. 10.1038/s41587-020-0435-1

Farid, M., & Navabi, A. (2015). N2 fixation ability of different dry bean genotypes. Canadian Journal of Plant Science, 95(6), 1243–1257. 10.4141/cjps-2015-084

Fréville, H., Montazeaud, G., Forst, E., David, J., Papa, R., & Tenaillon, M. I. (2022). Shift in beneficial interactions during crop evolution. Evolutionary Applications, 15(6), 905–918. 10.1111/eva.13390

Gaba, S., Lescourret, F., Boudsocq, S., Enjalbert, J., Hinsinger, P., Journet, E. P., Navas, M. L., Wery, J., Louarn, G., Malezieux, E., Pelzer, E., Prudent, M., & Ozier-Lafontaine, H. (2015). Multiple cropping systems as drivers for providing multiple ecosystem services : From concepts to design. Agronomy for Sustainable Development, 35(2), 607–623. 10.1007/s13593-014-0272-z

Girvan, M. S., Bullimore, J., Pretty, J. N., Osborn, A. M., & Ball, A. S. (2003). Soil type is the primary determinant of the composition of the total and active bacterial communities in arable soils. Applied and Environmental Microbiology, 69(3), 1800–1809. 10.1128/AEM.69.3.1800-1809.2003

Hauggaard-Nielsen, H., Jørnsgaard, B., Kinane, J., & Jensen, E. S. (2008). Grain legume–cereal intercropping : The practical application of diversity, competition and facilitation in arable and organic cropping systems. Renewable Agriculture and Food Systems, 23(1), 3–12. 10.1017/S1742170507002025

Houssard C. & Escarré J. (1991). The effects of seed weight on growth and competitive ability of *Rumex acetosella* from two successional old-fields. Oecologia, 86(2), 236–242. 10.1007/BF00317536.

Hupe, A., Naether, F., Haase, T., Bruns, C., Heß, J., Dyckmans, J., Joergensen, R. G., & Wichern, F. (2021). Evidence of considerable C and N transfer from peas to cereals via direct root contact but not via mycorrhiza. Scientific Reports, 11(1), 11424. 10.1038/s41598-021-90436-8

Johansen, A., & Jensen, E. S. (1996). Transfer of N and P from intact or decomposing roots of pea to barley interconnected by an arbuscular mycorrhizal fungus. Soil Biology & Biochemistry, 28(1), 73–81. 10.1016/0038-0717(95)00117-4

Laguerre, G., Courde, L., Nouaım, R., Lamy, I., Revellin, C., Breuil, M. C., & Chaussod, R. (2006). Response of rhizobial populations to moderate copper stress applied to an agricultural soil. Microbial Ecology, 52, 426–435. 10.1007/s00248-006-9081-5

Lauber, C. L., Strickland, M. S., Bradford, M. A., & Fierer, N. (2008). The influence of soil properties on the structure of bacterial and fungal communities across land-use types. Soil Biology and Biochemistry, 40(9), 2407–2415. 10.1016/j.soilbio.2008.05.021

Li, B., Li, Y.-Y., Wu, H.-M., Zhang, F.-F., Li, C.-J., Li, X.-X., Lambers, H., & Li, L. (2016a). Root exudates drive interspecific facilitation by enhancing nodulation and N2 fixation. Proceedings of the National Academy of Sciences of the United States of America, 113(23), 6496–6501. 10.1073/pnas.1523580113

Li, C., Dong, Y., Li, H., Shen, J., & Zhang, F. (2016b). Shift from complementarity to facilitation on P uptake by intercropped wheat neighboring with faba bean when available soil P is depleted. Scientific Reports, 66(1), 18663. 10.1038/srep18663

Li, C., Stomph, T.-J., Makowski, D., Li, H., Zhang, C., & Zhang, F. (2023). The productive performance of intercropping. Proceedings of the National Academy of Sciences of the United States of America, 120(2), e2201886120. 10.1073/pnas.2201886120

Li, H. (2021). Irrigation has a higher impact on soil bacterial abundance, diversity and composition than nitrogen fertilization. Scientific Reports, 11(1), 16901. 10.1038/s41598-021-96234-6

Li, H., Shen, J., Zhang, F., Clairotte, M., Drevon, J.-J., Le Cadre, E., & Hinsinger, P. (2008). Dynamics of phosphorus fractions in the rhizosphere of common bean (*Phaseolus vulgaris* L.) and durum wheat (*Triticum turgidum durum* L.) grown in monocropping and intercropping systems. Plant and Soil, 312(1), 139–150. 10.1007/s11104-007-9512-1

Li, L., Li, S. M., Sun, J. H., Zhou, L. L., Bao, X. G., Zhang, H. G., & Zhang, F. S. (2007). Diversity enhances agricultural productivity via rhizosphere phosphorus facilitation on phosphorus-deficient soils. Proceedings of the National Academy of Sciences of the United States of America, 104(27), 11192–11196. 10.1073/pnas.0704591104

Li, S. M., Li, L., Zhang, F. S., & Tang, C. (2004). Acid phosphatase role in chickpea/maize intercropping. Annals of Botany, 94(2), 297–303. 10.1093/aob/mch140

Loreau, M. & Hector A. (2001). Partitioning selection and complementarity in biodiversity experiments. Nature, 412, 72–76. 10.1038/35083573

Lun, A. T. L., Chen, Y., & Smyth, G. K. (2016). It’s DE-licious : a recipe for differential expression analyses of RNA-seq experiments using quasi-likelihood methods in edgeR. In E. Mathé & S. Davis (Eds.), Statistical Genomics (Vol. 1418, p. 391–416). Springer New York. 10.1007/978-1-4939-3578-9_19

Ma, H. (2023). Maize/alfalfa intercropping enhances yield and phosphorus acquisition. Field Crops Research, 303, 109136. 10.1016/j.fcr.2023.109136

Maitra, S., Hossain, A., Brestic, M., Skalicky, M., Ondrisik, P., Gitari, H., Brahmachari, K., Shankar, T., Bhadra, P., Palai, J. B., Jena, J., Bhattacharya, U., Duvvada, S. K., Lalichetti, S., & Sairam, M. (2021). Intercropping—A low input agricultural strategy for food and environmental security. Agronomy, 11(2), 343. 10.3390/agronomy11020343

Malviya, M. K., Solanki, M. K., Li, C.-N., Wang, Z., Zeng, Y., Verma, K. K., Singh, R. K., Singh, P., Huang, H.-R., Yang, L.-T., Song, X.-P., & Li, Y.-R. (2021). Sugarcane-legume intercropping can enrich the soil microbiome and plant growth. Frontiers in Sustainable Food Systems, 5, 606595. 10.3389/fsufs.2021.606595

Marmagne, A., Masclaux-Daubresse, C., & Chardon, F. (2022). Modulation of plant nitrogen remobilization and postflowering nitrogen uptake under environmental stresses. Journal of Plant Physiology, 277, 153781. 10.1016/j.jplph.2022.153781

Matsuoka, Y., Mitchell, S. E., Kresovich, S., Goodman, M., & Doebley, J. (2002). Microsatellites in *Zea* – variability, patterns of mutations, and use for evolutionary studies. Theoretical and Applied Genetics, 104(2), 436–450. 10.1007/s001220100694

Mazzola, M. (2004). Assessment and management of soil microbial community structure for disease suppression. Annual Review of Phytopathology, 42, 35–59. 10.1146/annurev.phyto.42.040803.140408

McMurdie, P. J., & Holmes, S. (2013). phyloseq : an R package for reproducible interactive analysis and graphics of microbiome census data. PLoS One, 8(4), e61217. 10.1371/journal.pone.0061217

Mead, R., & Willey, R. W. (1980). The concept of a ‘land equivalent ratio’ and advantages in yields from intercropping. Experimental Agriculture, 16(3), 217–228. 10.1017/S0014479700010978

Moyer-Henry, K. A., Burton, J. W., Israel, D. W., & Rufty, T. W. (2006). Nitrogen transfer between plants : A ^15^N natural abundance study with crop and weed species. Plant and Soil, 282(1), 7–20. 10.1007/s11104-005-3081-y

Nassary, E. K., Baijukya, F., & Ndakidemi, P. A. (2020). Productivity of intercropping with maize and common bean over five cropping seasons on smallholder farms of Tanzania. European Journal of Agronomy, 113, 125964. 10.1016/j.eja.2019.125964

Parnaudeau, V. V., Reau, R. R., & Dubrulle, P. P. (2012). Un outil d’évaluation des fuites d’azote vers l’environnement à l’échelle du système de culture : Le logiciel Syst’N. Innovations Agronomiques, 21, 59–70. http://www7.inra.fr/ciag/revue/volume_21_septembre_2012

Patel, Y., Zhu, C., Yamaguchi, T. N., Bugh, Y. Z., Tian, M., Holmes, A., Fitz-Gibbon, S. T., & Boutros, P. C. (2024). NFTest : Automated testing of Nextflow pipelines. Bioinformatics, 40(2), btae081. 10.1093/bioinformatics/btae081

Patro, R., Duggal, G., Love, M. I., Irizarry, R. A., & Kingsford, C. (2017). Salmon provides fast and bias-aware quantification of transcript expression. Nature Methods, 14(4), 417–419. 10.1038/nmeth.4197

Peña-Cabriales, J. J., & Castellanos, J. Z. (1993). Effects of water stress on N2 fixation and grain yield of *Phaseolus vulgaris* L. Plant and Soil, 152(1), 151–155. 10.1007/BF00016345

Peoples, M. B., Boddey, R. M., & Herridge, D. F. (2002). Quantification of nitrogen fixation (Chapter 13). In G. J. Leigh (Ed.), Nitrogen Fixation at the Millennium (p. 357–389). Elsevier Science. 10.1016/B978-044450965-9/50013-6

Poggio, S. L. (2005). Structure of weed communities occurring in monoculture and intercropping of field pea and barley. Agriculture, Ecosystems & Environment, 109(1), 48–58. 10.1016/j.agee.2005.02.019

Ritchie, M. E., Phipson, B., Wu, D., Hu, Y., Law, C. W., Shi, W., & Smyth, G. K. (2015). Limma powers differential expression analyses for RNA-sequencing and microarray studies. Nucleic Acids Research, 43(7), e47. 10.1093/nar/gkv007

Robinson, M. D., & Oshlack, A. (2010). A scaling normalization method for differential expression analysis of RNA-seq data. Genome Biology, 11(3), R25. 10.1186/gb-2010-11-3-r25

Rodriguez, C., Carlsson, G., Englund, J.-E., Flöhr, A., Pelzer, E., Jeuffroy, M.-H., Makowski, D., & Jensen, E. S. (2020). Grain legume-cereal intercropping enhances the use of soil-derived and biologically fixed nitrogen in temperate agroecosystems. A meta-analysis. European Journal of Agronomy, 118, 126077. 10.1016/j.eja.2020.126077

Rokhbakhsh-Zamin, F., Sachdev, D., Kazemi-Pour, N., Engineer, A., Pardesi, K. R., Zinjarde, S. S., Dhakephalkar, P. K., & Chopade, B. A. (2011). Characterization of plant-growth-promoting traits of *Acinetobacter* species isolated from rhizosphere of *Pennisetum glaucum*. Journal of Microbiology and Biotechnology, 21(6), 556–566. 10.4014/jmb.1012.12006

Schlatter, D. C., Bakker, M. G., Bradeen, J. M., & Kinkel, L. L. (2015). Plant community richness and microbial interactions structure bacterial communities in soil. Ecology, 96(1), 134–142. 10.1890/13-1648.1

Shearer, G., & Kohl, D. H. (1988). Natural ^15^N abundance as a method of estimating the contribution of biologically fixed nitrogen to N2-fixing systems : Potential for non-legumes. Plant and Soil, 110(2), 317–327. 10.1007/BF02226812

Shearer, G., Kohl, D. H., & Harper, J. E. (1980). Distribution of ^15^N among plant parts of nodulating and nonnodulating isolines of soybeans. Plant Physiology, 66(1), 57–60. 10.1104/pp.66.1.57

Smith, T., Heger, A., & Sudbery, I. (2017). UMI-tools : Modeling sequencing errors in Unique Molecular Identifiers to improve quantification accuracy. Genome Research, 27(3), 491–499. 10.1101/gr.209601.116

Ulbrich, T. C., Rivas-Ubach, A., Tiemann, L. K., Friesen, M. L., & Evans, S. E. (2022). Plant root exudates and rhizosphere bacterial communities shift with neighbor context. Soil Biology & Biochemistry, 172, 108753. 10.1016/j.soilbio.2022.108753

Vazeux-Blumental, N., Manicacci, D., & Tenaillon, M. I. (2024). The milpa, from Mesoamerica to present days, a traditional agricultural system serving agroecology. Comptes rendus de l’Académie des Sciences Biologies, 6(347), 159–173. 10.5802/crbiol.164

Voisin, A.-S., Salon, C., Munier-Jolain, N. G., & Ney, B. (2002). Effect of mineral nitrogen on nitrogen nutrition and biomass partitioning between the shoot and roots of pea (*Pisum sativum* L.). Plant and Soil, 242(2), 251–262. 10.1023/A:1016214223900

Xing, Y., Yu, R.-P., An, R., Yang, N., Wu, J.-P., Ma, H.-Y., Zhang, J.-D., Bao, X.-G., Lambers, H., & Li, L. (2023). Two pathways drive enhanced nitrogen acquisition via a complementarity effect in long-term intercropping. Field Crops Research, 293, 108854. 10.1016/j.fcr.2023.108854

Yan, J., Han, X. Z., Ji, Z. J., Li, Y., Wang, E. T., Xie, Z. H., & Chen, W. F. (2014). Abundance and diversity of soybean-nodulating rhizobia in black soil are impacted by land use and crop management. Applied and Environmental Microbiology, 80(17), 5394–5402. 10.1128/AEM.01135-14

Yin, C., Casa Vargas, J. M., Schlatter, D. C., Hagerty, C. H., Hulbert, S. H., & Paulitz, T. C. (2021). Rhizosphere community selection reveals bacteria associated with reduced root disease. Microbiome, 9(1), 86. 10.1186/s40168-020-00997-5

Zeng, J., Liu, X., Song, L., Lin, X., Zhang, H., Shen, C., & Chu, H. (2016). Nitrogen fertilization directly affects soil bacterial diversity and indirectly affects bacterial community composition. Soil Biology & Biochemistry, 92, 41–49. 10.1016/j.soilbio.2015.09.018

Zhang, C., Postma, J. A., York, L. M., & Lynch, J. P. (2014). Root foraging elicits niche complementarity-dependent yield advantage in the ancient ‘three sisters’ (maize/bean/squash) polyculture. Annals of Botany, 114(8), 1719–1733. 10.1093/aob/mcu191

Zhang, D., Zhang, C., Tang, X., Li, H., Zhang, F., Rengel, Z., Whalley, W. R., Davies, W. J., & Shen, J. (2016). Increased soil phosphorus availability induced by faba bean root exudation stimulates root growth and phosphorus uptake in neighbouring maize. New Phytologist, 209(2), 823–831. 10.1111/nph.13613

Zhang, S., Lehmann, A., Zheng, W., You, Z., & Rillig, M. C. (2019). Arbuscular mycorrhizal fungi increase grain yields : A meta-analysis. New Phytologist, 222(1), 543–555. 10.1111/nph.15570

Zizumbo-Villarreal, D., & Colunga-GarcíaMarín, P. (2010). Origin of agriculture and plant domestication in West Mesoamerica. Genetic Resources and Crop Evolution, 57(6), 813–825. 10.1007/s10722-009-9521-4

