## Supplemental Figures for "Revival of traditional agricultural systems – A multidisciplinary on-farm survey of maize-bean intercropping reveals unexpected competition effects on beans"

### *Plants, People, Planet* Supporting Information

Article title: A multidisciplinary on-farm survey of conventional maize-bean intercropping reveals unexpected competition effects on bean yield.

Authors: Noa Vazeux-Blumental, Laura Mathieu, Théo Trabac, Carine Palaffre, Bernard Lagardère, Maryse Carraretto, Cyril Bauland, Martine Le Guilloux, Christine Paysant Le Roux, José Caïus, Anne Marmagne, Jérôme Enjalbert, Timothée Flutre, Edith Le Cadre, Virginie Parnaudeau, Daniel Muller, Yvan Moënne-Loccoz, Domenica Manicacci, Maud Tenaillon.

The following Supporting Information is available for this article:

**Figure S1** Geographical location of the seven farms F1 to F7 (left) near Tarbes (right) in southwestern France.

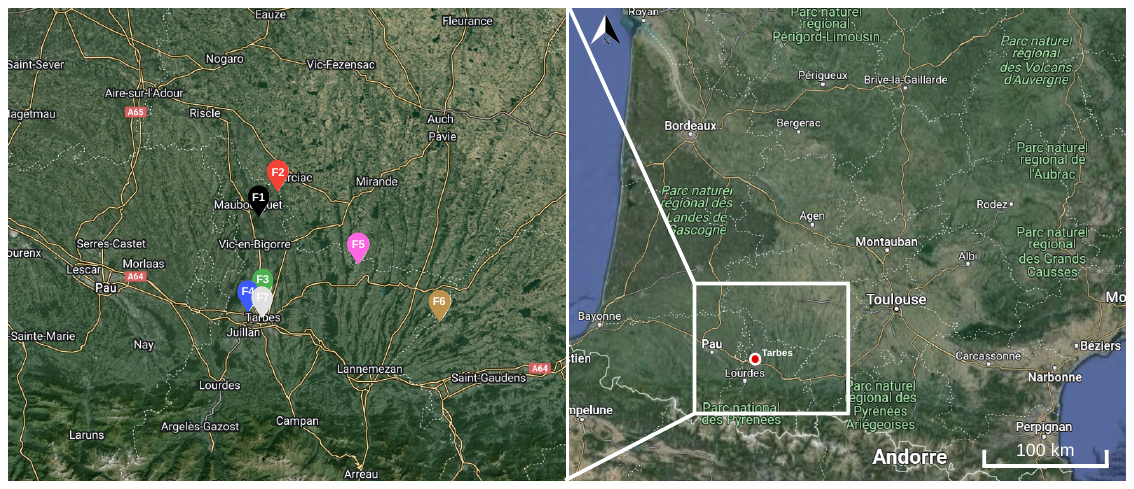

**Figure S2** Boxplot of the 21 R-squared values between blocks calculated from RNSAseq gene counts. The two dots with the lowest values indicate the correlations calculated for the bean samples in the modality M2B between block 1 and block 2 (B1_M2B_1.2) and between block 1 and block 3 (B_M2B_1.3).

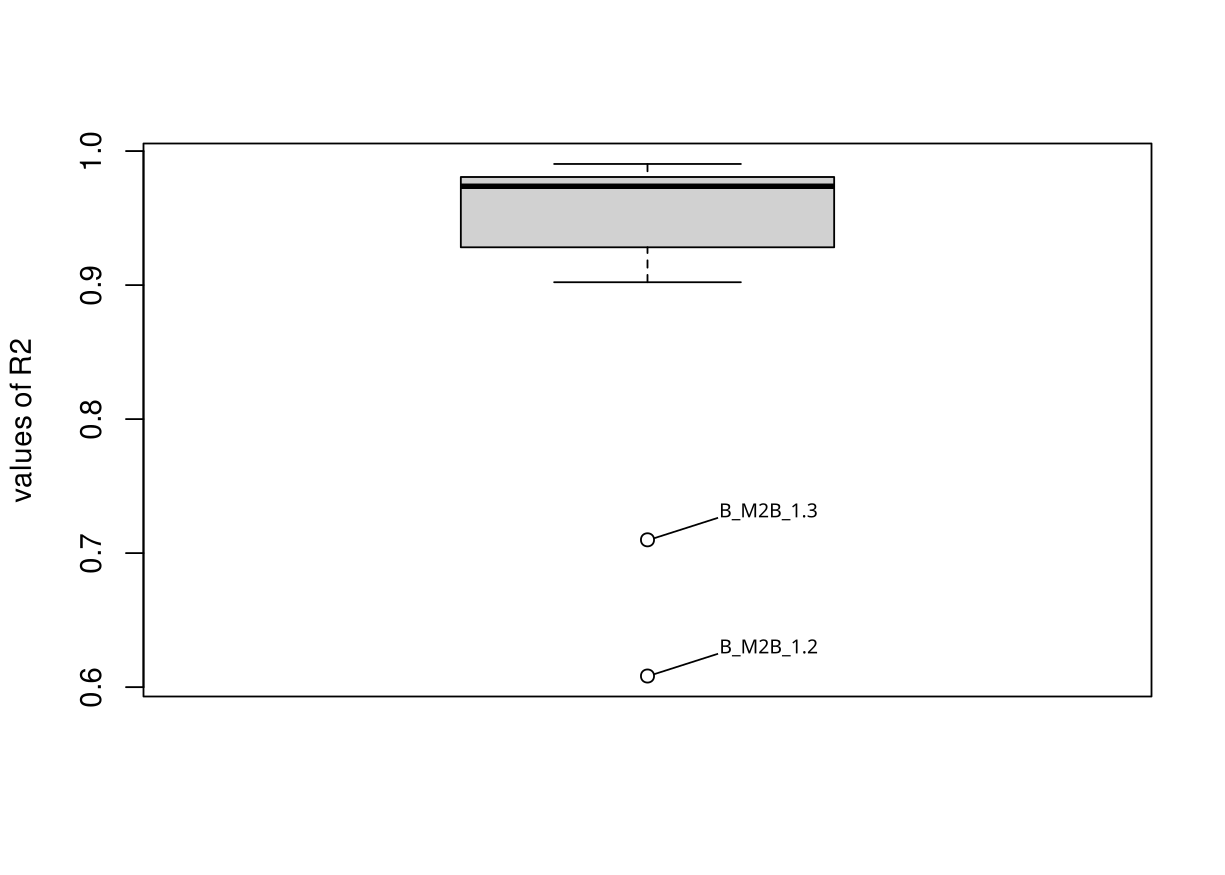

**Figure S3** Soil texture of the fields studied in South-West France, based on their composition in clay (Clay), silt (Fsi, Csi) and sand (Csa, Fsa), represented according to the farm (F1 to F7) and cropping system (B = Bean, M = Maize, MB = Intercropping of maize and bean).

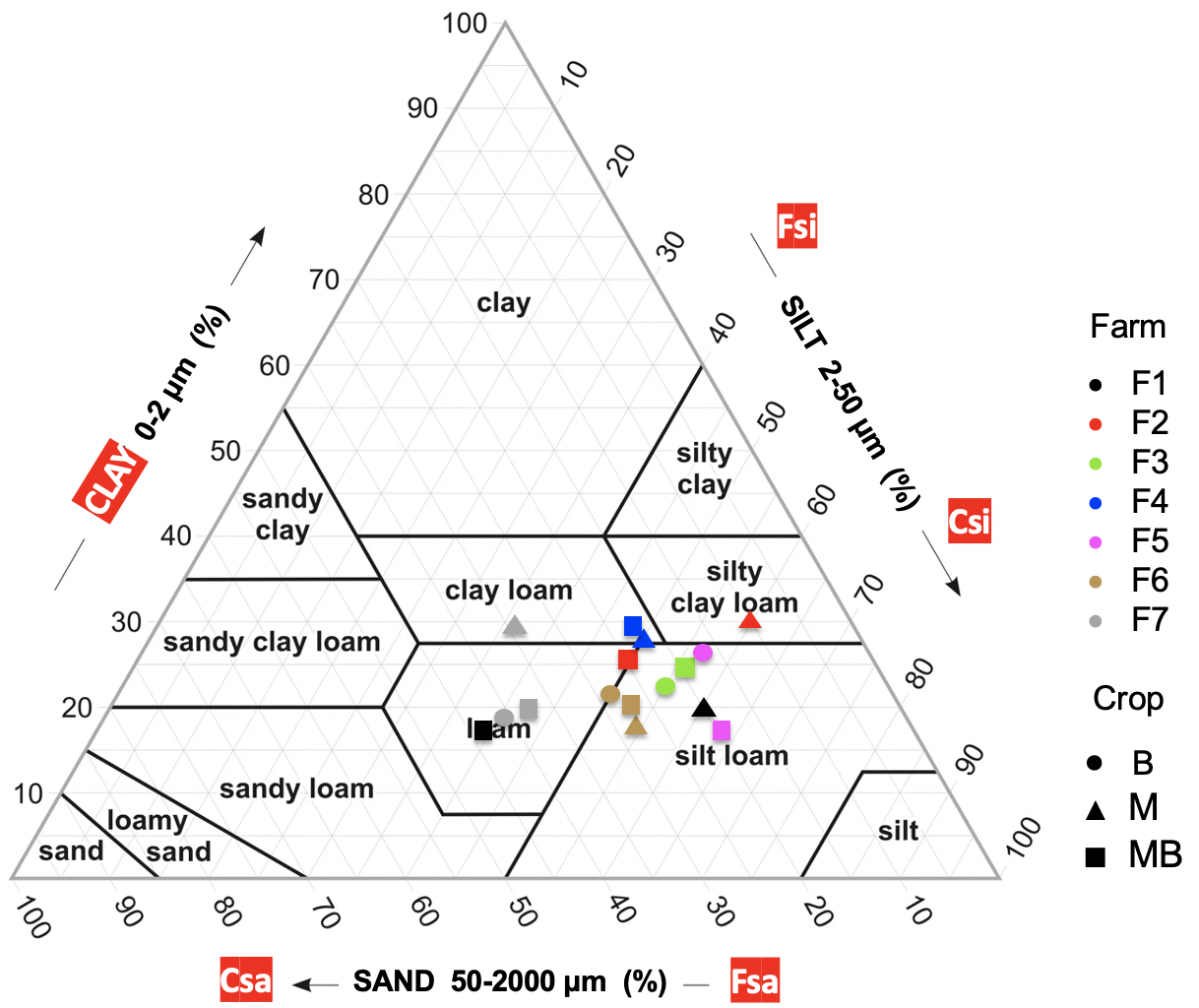

**Figure S4** Mean relative abundance of the 10 most abundant taxa at various taxonomic levels (phylum, class, order, family, genus and species) for bulk and rhizospheric soils of sole-cropped bean, sole-cropped maize, intercropped bean and intercropped maize.

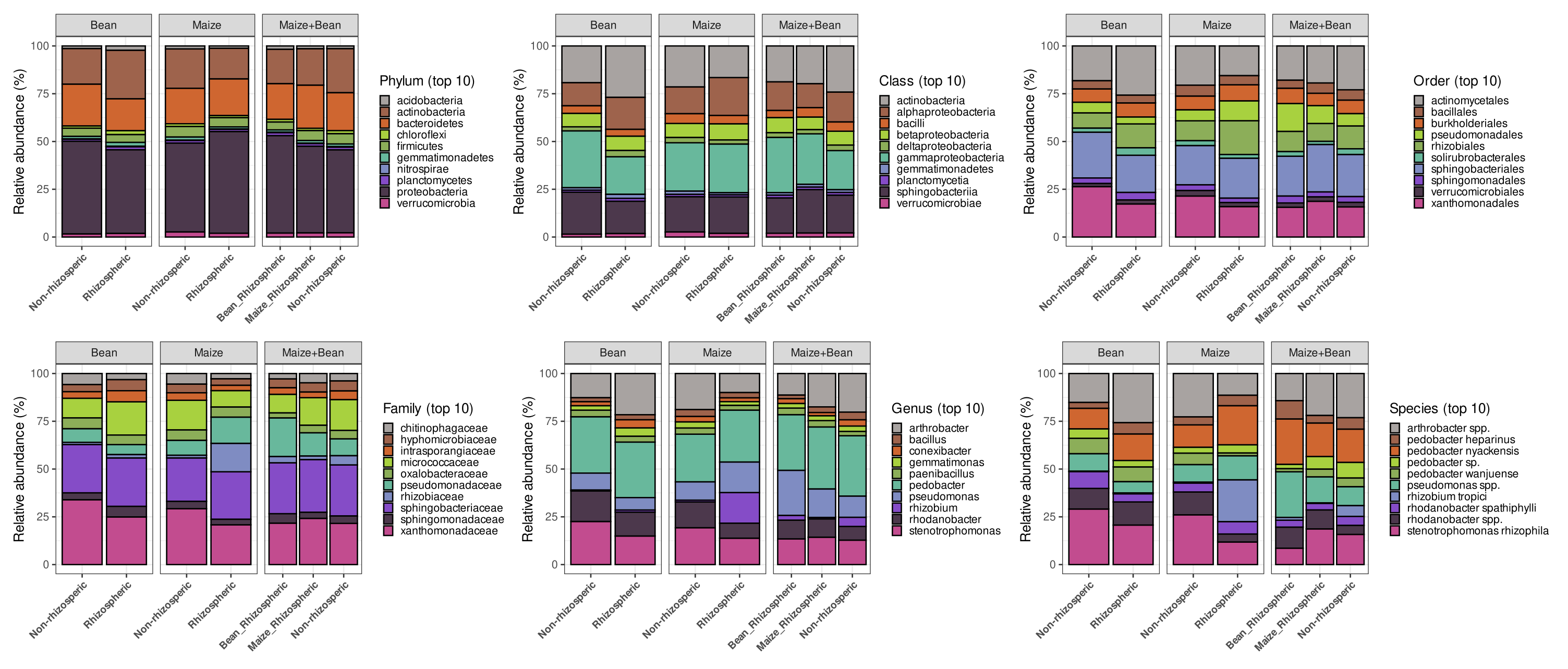

**Figure S5** Alpha-diversity of soil bacterial communities in different cropping systems.
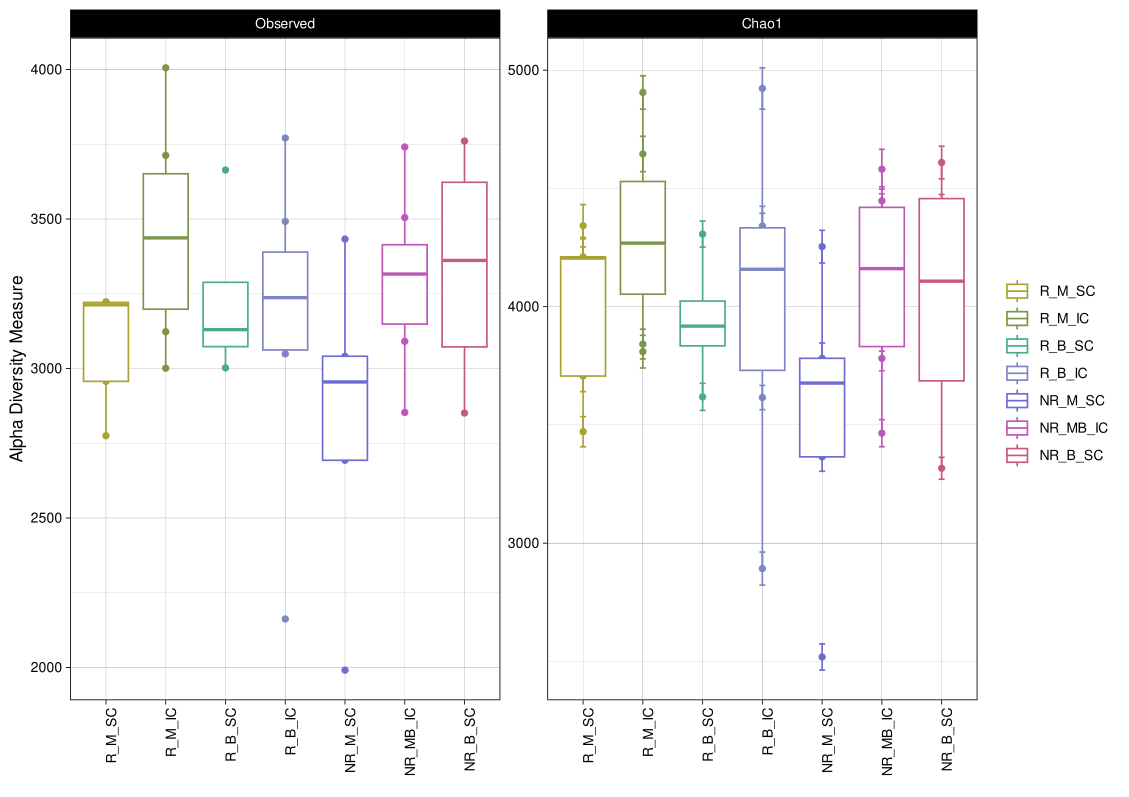
Boxplots with dots of observed OTUs and Chao1 index are shown for rhizospheric and non-rhizospheric soils of sole-cropped beans (respectively R_B_SC and NR_B_SC), rhizospheric and non-rhizospheric soils of sole-cropped maize (R_M_SC and NR_M_SC), rhizospheric soil of intercropped beans and maize (R_B_IC and R_M_IC), and non-rhizospheric soil of intercropped beans and maize (NR_MB_IC).

**Figure S6** Non-metric Multi Dimensional Scaling (NMDS) representation of farm samples using bacterial metabarcoding data. The NMDS was generated from Unifrac distances calculated for 11,586 16S rRNA-based OTUs across different modalities: rhizospheric and non-rhizospheric soils of sole cropped beans (respectively R_B_SC and NR_B_SC), rhizospheric and non-rhizospheric soils of sole cropped maize (R_M_SC and NR_M_SC), rhizospheric soils of intercropped beans and maize (R_B_IC and R_M_IC), and non-rhizospheric soils of intercropped beans and maize (NR_MB_IC). The shape of points represents the modality, while the color refers to the corresponding farm.

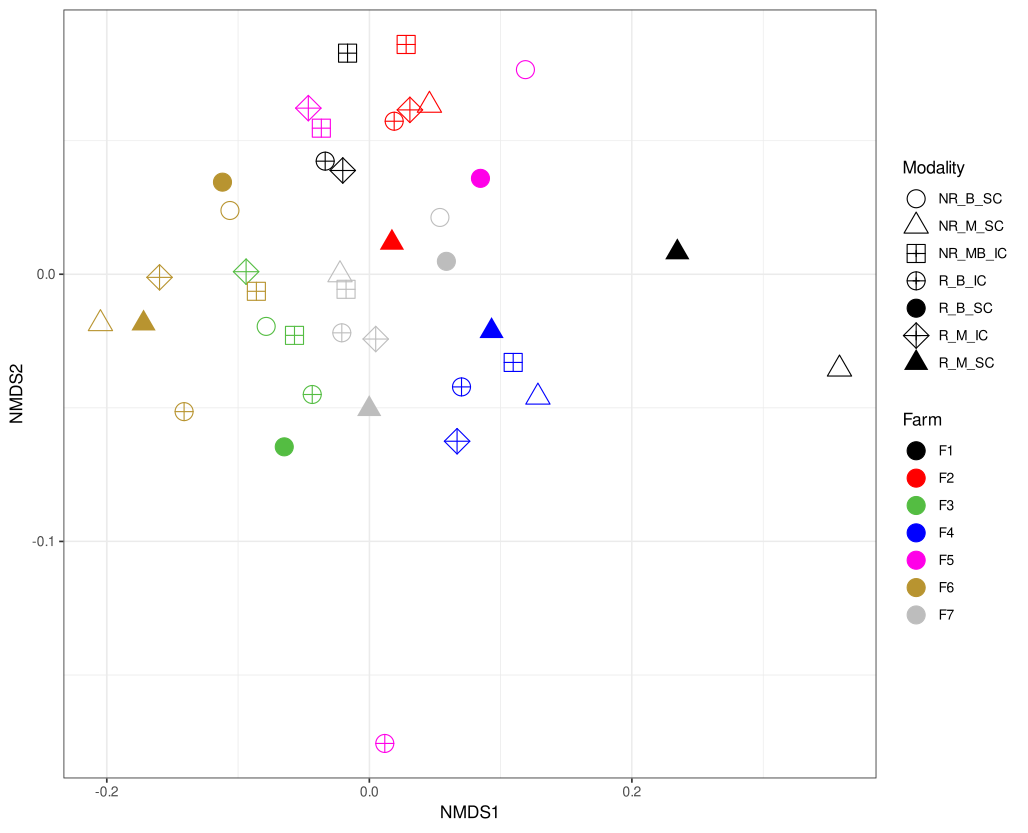

**Figure S7** Co-inertia analysis between (i) 13 soil variables, irrigation level, simulated soil nitrogen mineralization (Nsim), nitrogen units (Nin) supplied during the growing season, and (ii) bacterial OTUs in soil samples. Farms F1 to F7 are indicated. The co-inertia coefficient RV between the two datasets was 0.58 (*P* value = 0.025), obtained from 10,000 permutations in Monte Carlo simulations. Yellow dots represent soil profiles, while blue dots represent the OTU profile of the farm. Shorter line lengths indicate greater consistency between the two datasets.

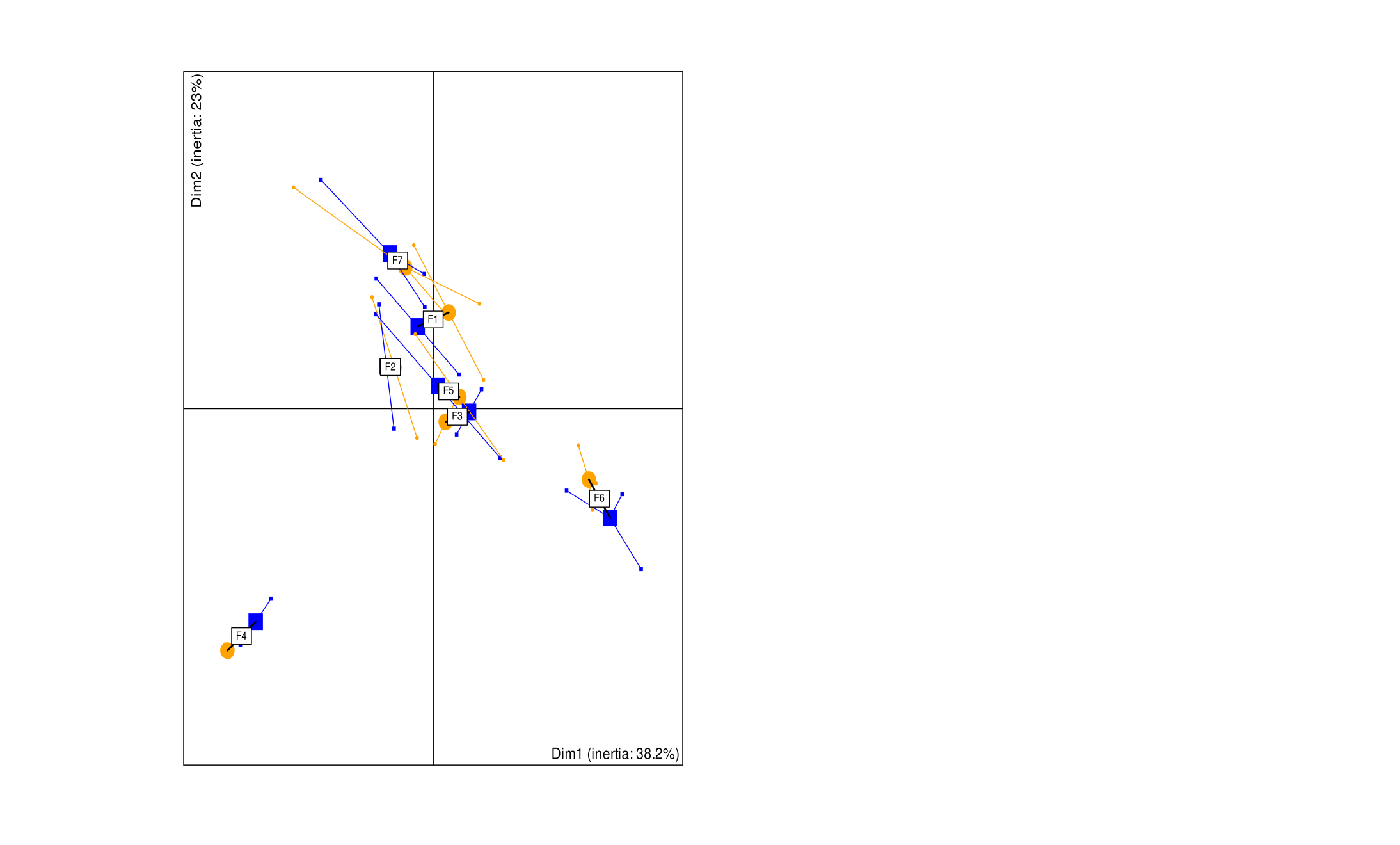

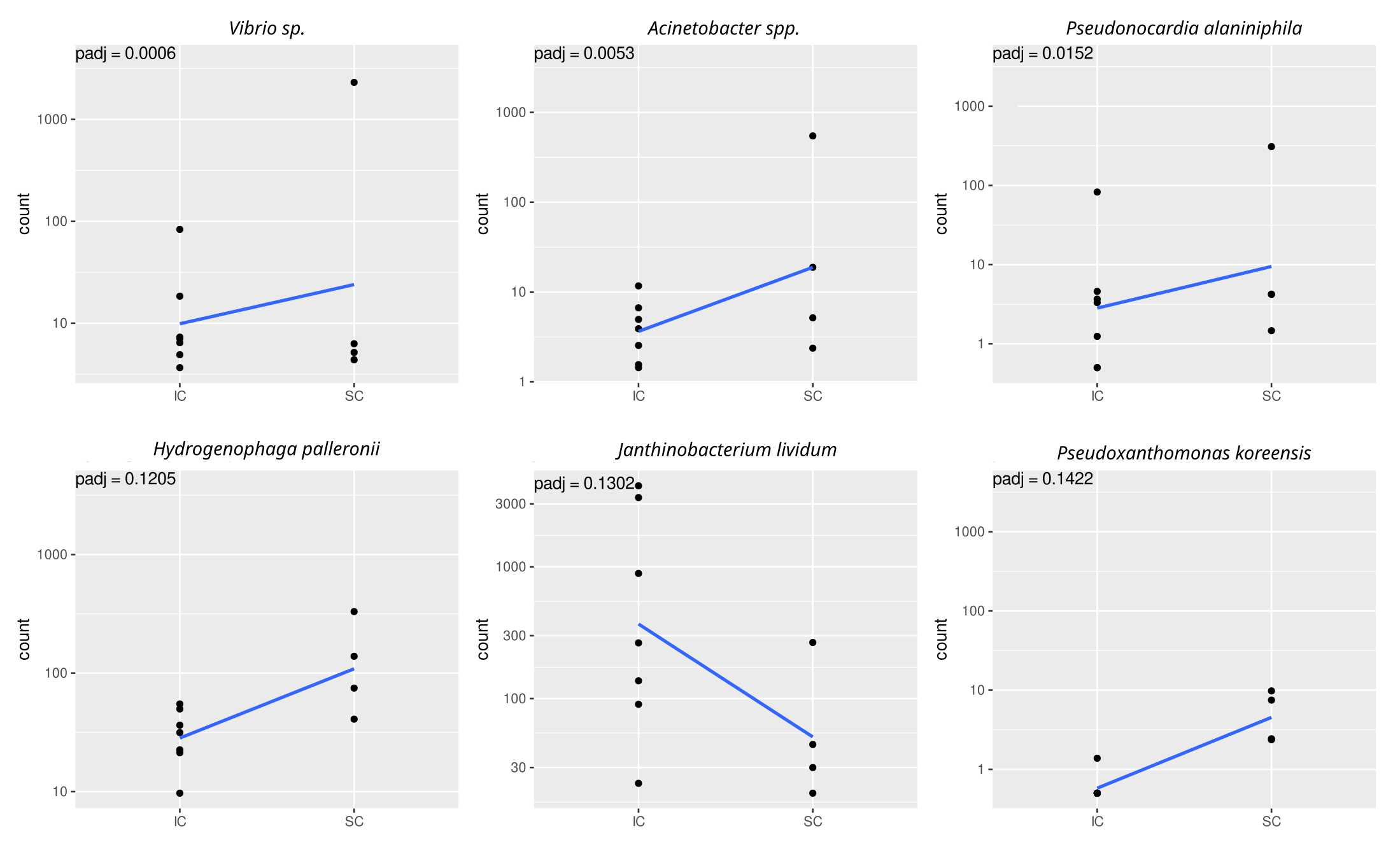
**Figure S8** Discriminant bacterial OTUs of bean rhizosphere in the two cropping systems, i.e. intercropped bean (IC) vs sole-cropped bean (SC). The six OTUs with the lowest FDR adjusted *P* values are shown.
**Figure S9** Discriminant bacterial OTUs of maize rhizosphere between the two cropping systems, i.e. intercropped maize (IC) vs sole-cropped maize (SC). The six OTUs with the lowest FDR adjusted *P* values are shown. Bacterium SOSP1-30 is now affiliated to *Ktedonobacter robiniae*.

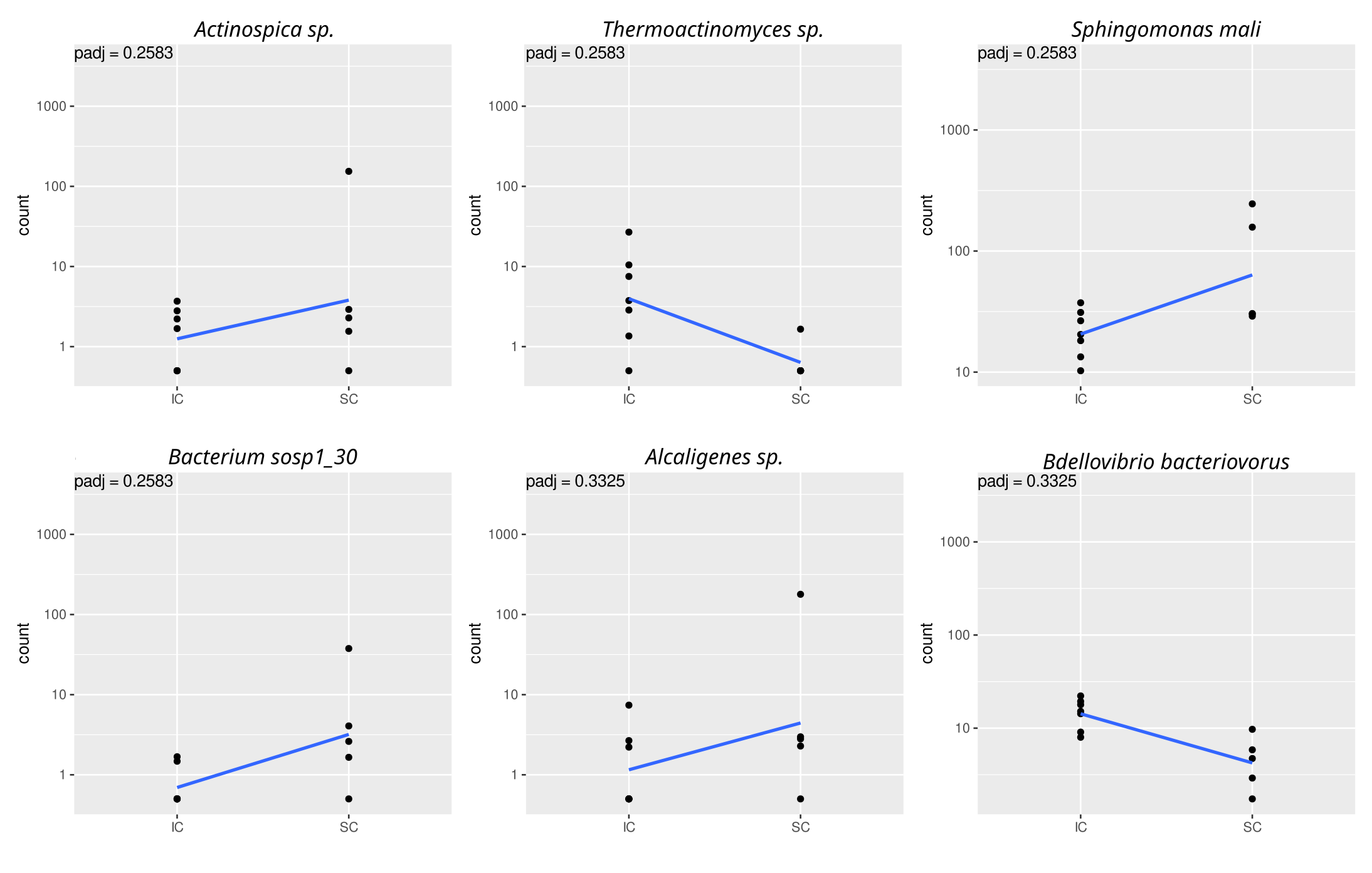

**Figure S10** Plots for maize phenotypic traits displaying significant Farm × Cropping system interaction. Mean values per farm are given for each cropping system, i.e. sole-cropped maize (red) and intercropped maize (black). The complete phenotypic trait names are given in Table 1. Phenotypic traits (and their units in parentheses) are indicated above each panel, along with significant effects in the two-way ANOVA (F = Farm effect, Tr = Cropping system effect, F*Tr = Farm × Cropping system interaction).

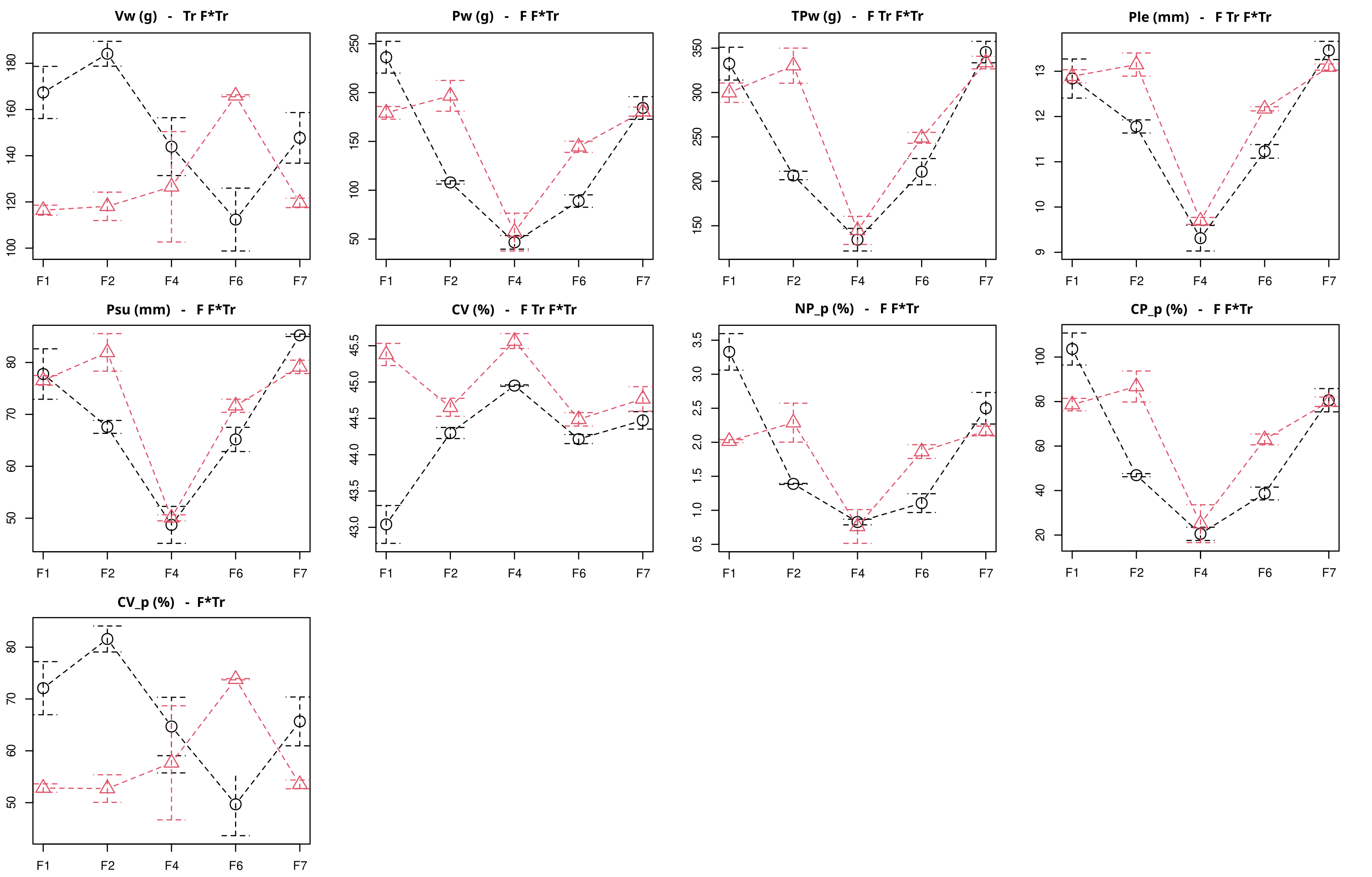

**Figure S11** Plots for bean phenotypic traits displaying significant Farm × Cropping system interaction. Mean values per farm are given for each cropping system, i.e. sole-cropped bean (red) and intercropped bean (black). The complete phenotypic trait names are given in Table 1. Phenotypic traits (and their units in parentheses) are indicated above each panel, along with significant effects in the two-way ANOVA (F = Farm effect, Tr = Cropping system effect, F*Tr = Farm × Cropping system interaction).

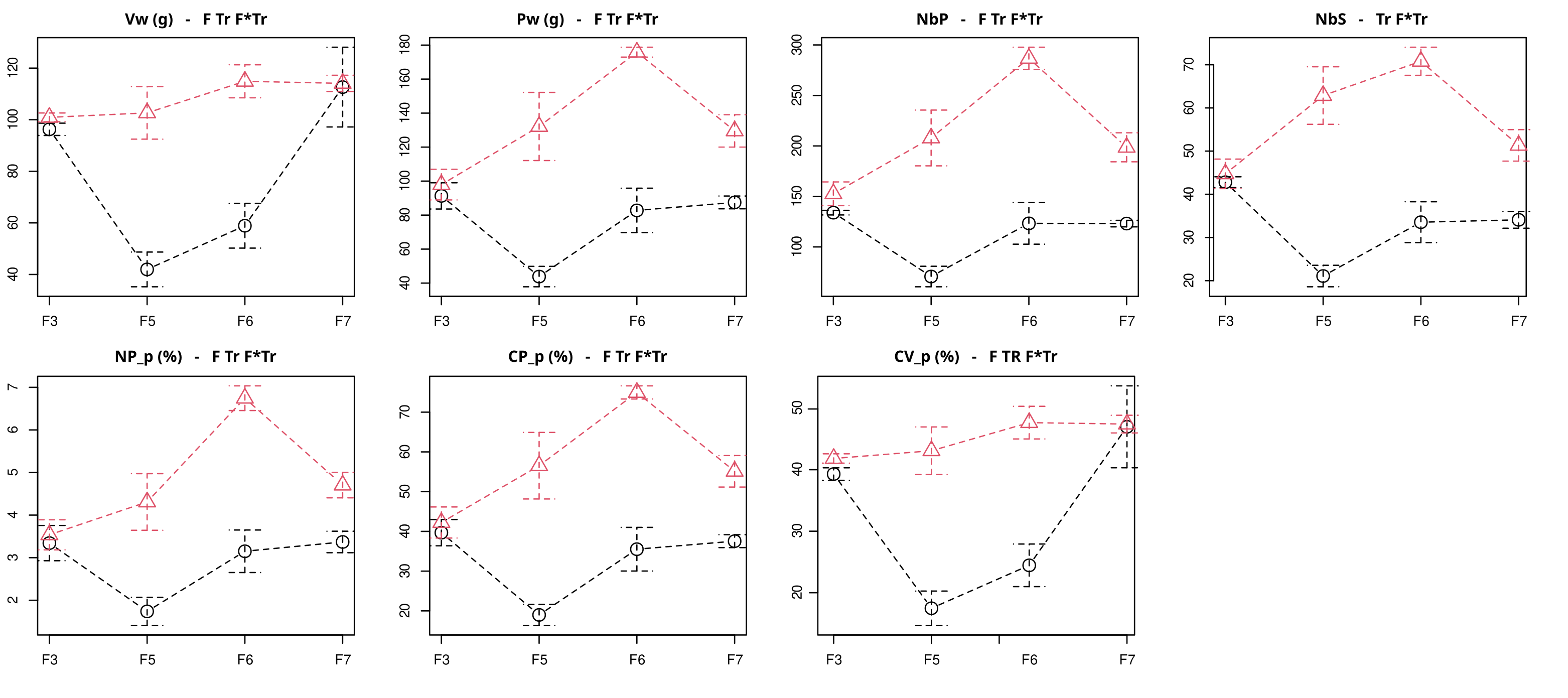

**Figure S12** Correlation graph between maize and bean intercropped plants. Rows refer to traits measured on bean and columns to traits measured on maize. Full trait names are indicated in Table 1. Ellipses show correlations for *Q* values < 0.2 and color shades indicate Pearson’s correlation values.

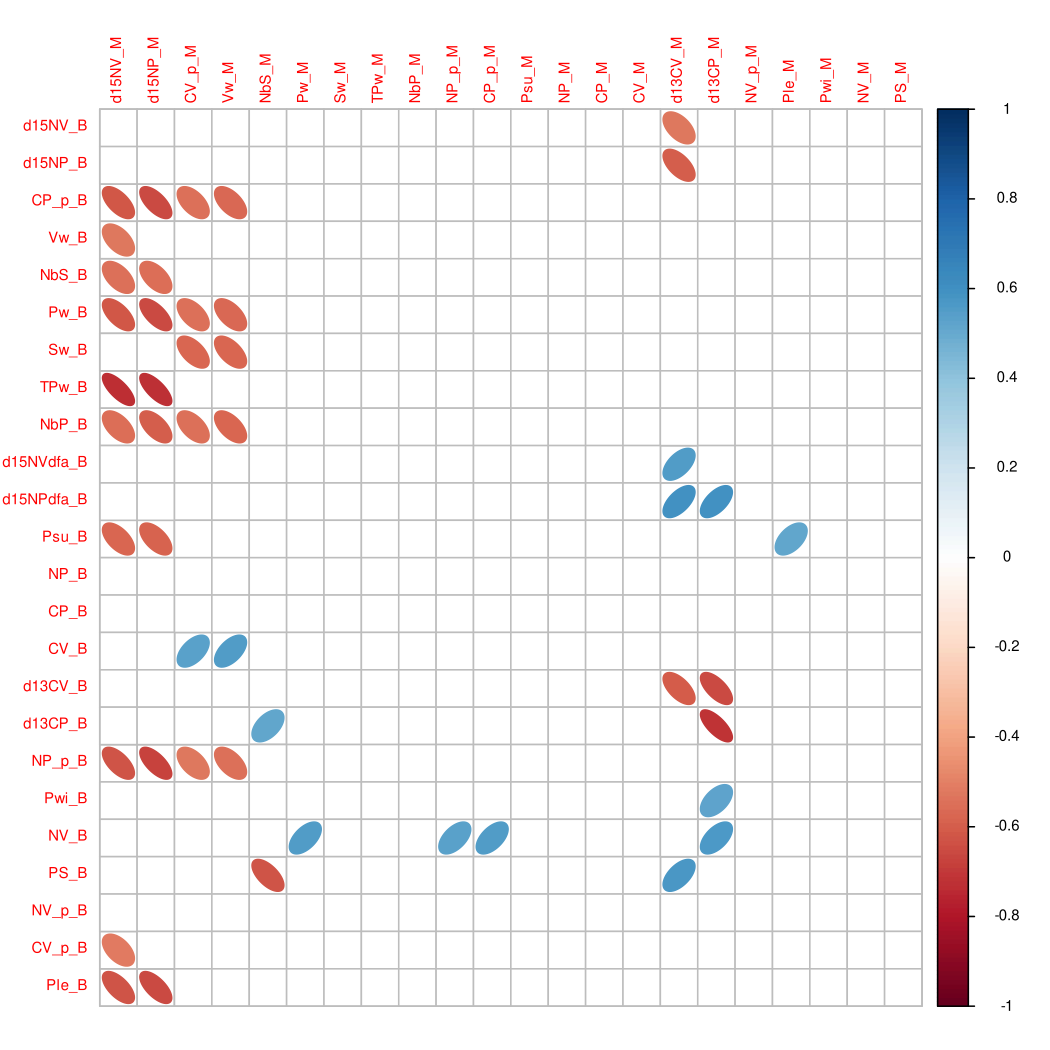

**Figure S13** Stripcharts of bean traits displaying significant effect of the cropping condition (bean sole cropping vs intercropped bean, *P* value < 0.05). **A**: width of the main leaflet, **B**: foliage density, **C**: number of seeds per plant, and D: number of pods per plant. Blue points represent the three values per cropping condition, and the horizontal lines indicate the mean for the corresponding cropping system i.e. sole-cropped bean (B), bean intercropped with maize genotype 1 (M1B), and bean intercropped with maize genotype 2 (M2B).

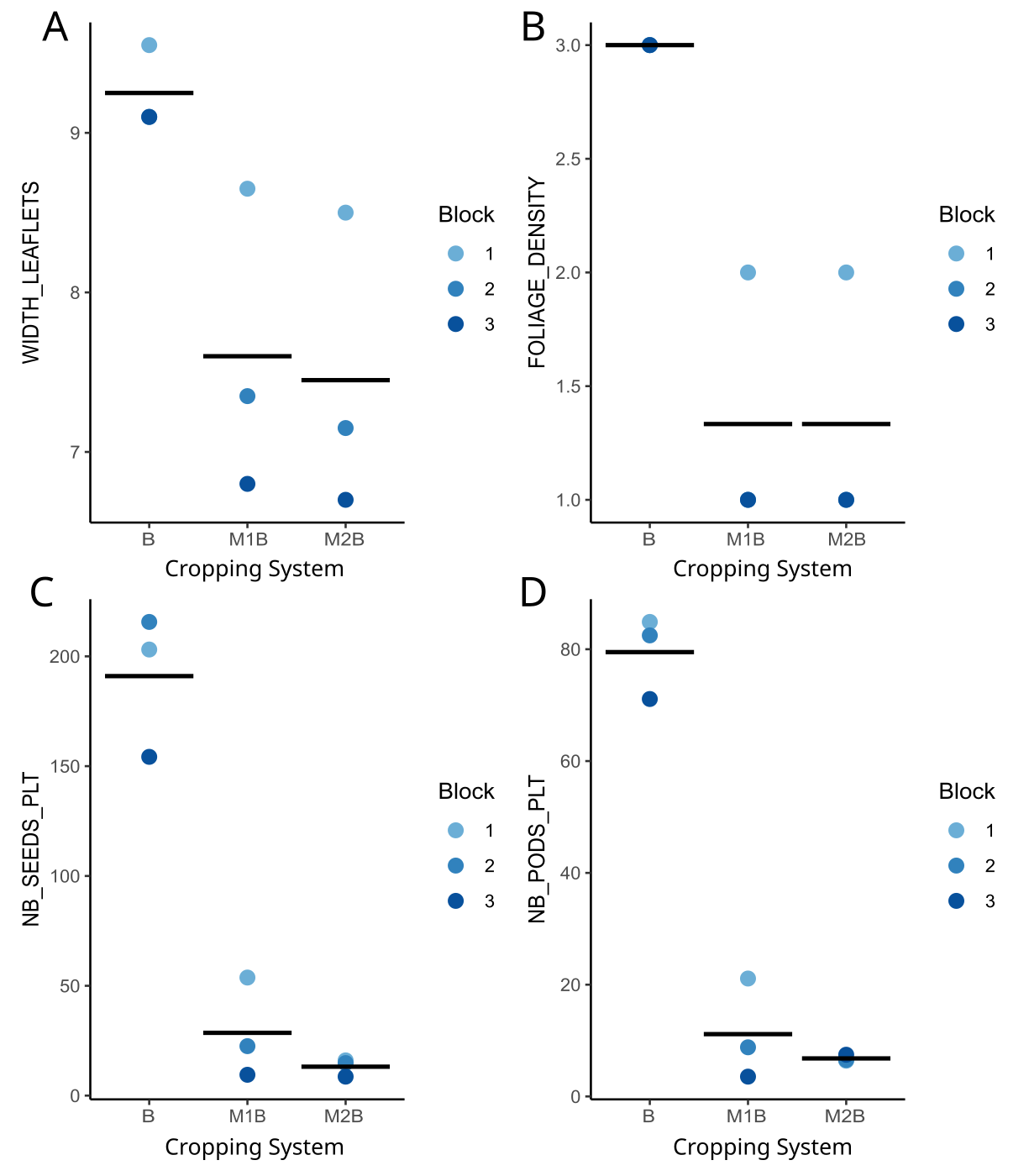

**Figure S14** Stripcharts of the maize traits displaying significant maize genotypic effect (*P* value < 0.05). **A**: number of days to silk emergence, **B**: one-thousand kernel weight, **C**: number of seeds per plant, and **D**: number of lodged plant in the row. Blue points represent the three values per cropping condition, and the horizontal lines indicate the mean for the corresponding cropping system i.e. sole-cropped maize genotype 1 (M1), sole-cropped maize genotype 2 (M2), intercropped maize genotype 1 (M1B), and intercropped maize genotype 2 (M2B).

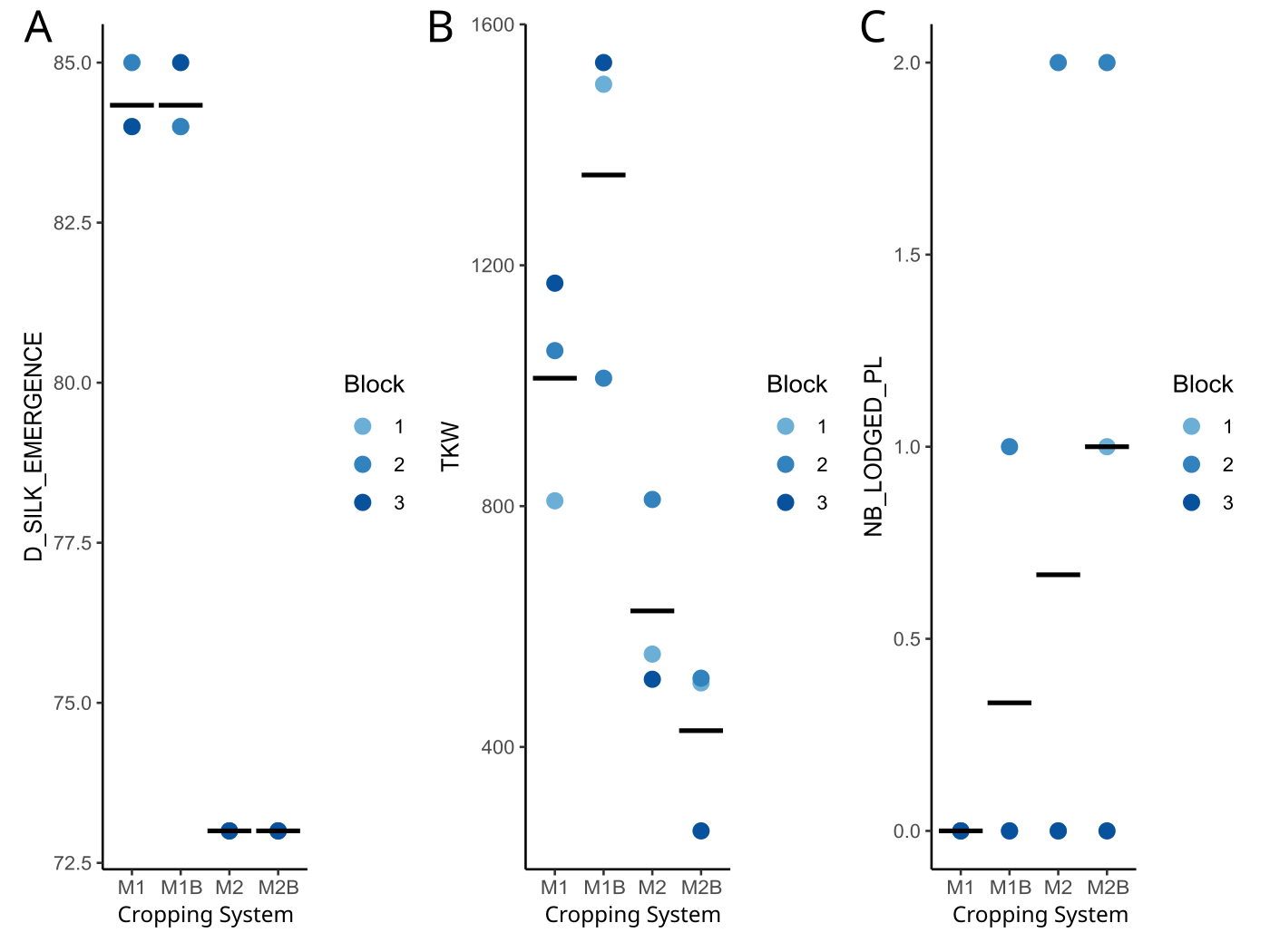

**Dataset S1** Description of fields and practices.

Cropping system refers to Sole Cropping (SC) of maize or beans, or Intercropping (IC) of maize and beans. Density for maize and bean corresponds to the number of plants per square meter, and Nin indicates the number of nitrogen units per hectare.

**Dataset S2** Description of variables measured on non-rhizospheric soil asured on samples pooled from the 3 plots of each field.

Ni and Am were measured on each of the three samples of soil per field, while all other variables were measured on pools of 3 samples, one taken from the 3 plots of each field.

**Dataset S3** Values of 13 soil variables and soil nitrogen mineralization.

Non-rhizospheric soil was sampled from 3 replicates (Rep) per farm per treatment. These replicates correspond to the ones used to collect the plants. Except for Ni and Am, soil variables were measured after pooling the 3 replicates (bulk). The last column (Nsim) refers to the values obtained from the simulations of soil nitrogen mineralization.

**Dataset S4** Data for 23 phenotypic and nutrition variables.

Measures were taken from 3 replicates (Rep) per field. Each replicate encompassed 3 to 5 plants (Plant). Variables are described in Table 2.

**Dataset S5** OTUs counts in 39 rhizospheric (R) and non-rhizospheric (NR) soil samples.

OTUs counts were determined after pooling samples of soil collected from n plants (n=Plants) from 3 replicates (Rep). The replicates correspond to the ones used to collect the plants (Table S4) and soil data (Table S3).

**Dataset S6** Description of OTUs.

**Dataset S7** RNAseq report of the 21 libraries.

Names and codes of libraries are provided along with details of the samples (Bean or Maize - M1, M2 genotypes, cropping system and block) and the number of reads before and after filtering.

**Dataset S8** Description of plant variables measured on the experimental assay in Saclay.

**Dataset S9** Significance of farm and cropping system factors in the ANOVA for the 13 soil variables, simulation of soil nitrogen mineralization (Nsim), nitrogen unite per hectare (Nin) and irrigation.

Description of soil variables are given in Table2.

**Dataset S10** Pearson's correlation coefficients and associated P-values (P) between Nsim and sol variables (only significant correlations are reported).

Soil variable definitions are given Table S2. *: P < 0.05, ** : P < 0.01, and ***: P < 0.001.

**Dataset S11** Effects of farm, rhizospheric compartment, cropping system and species on OTUs composition among sampling tested by PERMANOVA on unifrac distances.

**Dataset S12** Canonical weights of soil variables and practice variables to the coinertia axes

**Dataset S13** Significance level (p-values) of farm, cropping system and interaction on agronomic traits for maize and bean.

Variables definitions are given Table S2. *: P < 0.05, ** : P < 0.01, and ***: P < 0.001.
